# Z-DNA formation induces the totipotent-like state and primes Zscan4-dependent chromatin compartmentalization

**DOI:** 10.1101/2025.03.18.643869

**Authors:** Shireen Shajahan, Yann Loe-Mie, Marion Salmon-Legagneur, Tatiana Traboulsi, Agnès Thierry, Anne Dejean, Jack-Christophe Cossec

## Abstract

A remarkable transition during murine development is the progression from the 1-cell embryo to the 2-cell stage, accompanied by the activation of a specific set of embryonic genes, epigenome reprogramming, and nuclear architecture reorganization. Some of these characteristics are recapitulated *in vitro* with the spontaneous emergence of 2-cell-like cells from mouse embryonic stem cells, which exhibit a transcriptomic signature resembling the 2-cell stage, including the expression of genes such as *Dux*, *Zscan4*, and the repetitive element *MERVL*, and a more relaxed chromatin state. Here, we show that inter- and intra-chromosomal interactions, driven by Zscan4 chromatin factors, form during this transition and segregate into a distinct genomic compartment (Z compartment). Mechanistically, the formation of Z-DNA, an alternative DNA conformation regulated by polyamine levels, promotes the emergence of totipotent-like cells and the establishment of the Z compartment. This compartment is characterized by a decrease in active histone marks and a reduced expression of genes associated with differentiation and late developmental processes. Overall, these findings suggest that Z-DNA formation may play a dual role, first in activating ZGA genes and later in guiding genome compartmentalization to safeguard the totipotent-like state by restricting the expression of non-ZGA genes within a permissive chromatin environment.

## Main

Genome organization plays a crucial role in controlling gene expression and overall genome function by spatially arranging chromatin and its regulatory elements within the nucleus ^1–4^. A fundamental aspect of this organization is the division of the genome into two main compartments: compartment A, which is gene-rich and transcriptionally active, and compartment B, which is gene-poor and transcriptionally inactive ^5^. Building on this, the genome is further subdivided into Topologically Associating Domains (TADs) ^6–8^. These are regions where chromatin interactions are more frequent within the domain than with neighboring regions, allowing for localized regulation of gene expression. TADs ensure that regulatory elements like enhancers interact with the correct gene promoters, forming chromatin loops that maintain precise control over gene activation. In addition to TADs, Lamin-Associated Domains (LADs) contribute to genome organization by anchoring portions of the genome enriched for H3K9me2/3 to the nuclear periphery ^9^. These regions are typically transcriptionally repressed, adding another layer of spatial control by segregating inactive genes. Together, these structural features create a dynamic 3-dimensional (3D) genome architecture that is essential for controlling gene expression, maintaining genome stability, and guiding cellular identity during development and differentiation.

The emergence of totipotency at the zygote stage after the fertilization of two fully differentiated parental gametes is marked by a notable spatial reorganization of the paternal and maternal genomes, as well as the associated epigenomes ^10,11^. In the mouse zygote, the overall genome organization is loosely structured with weak A/B compartmentalization and poorly defined TAD organization ^12–14^. The zygotic histone marks are either inherited from the gametes, such as the non-canonical maternal patterns of H3K4me3, H3K27me3, and H2AK119ub1, or established *de novo*, such as H3K27ac, or erased and then re-established in a non-canonical way, such as H3K9me3 ^15–19^. These dynamics are associated with the establishment of specific genomic compartmentalization, such as the formation of LADs at the zygote stage, which may depend on the remodeling of H3K4 methylation but not on H3K9me2/3 as is usually the case ^20^, and of polycomb-associated domains (PADs), enriched in non-canonical H3K27me3 patterns observed in early 2-cell (2C) embryos ^21^. During mouse development, the parental-to-zygotic transition leads to the transcriptional activation of the zygotic genome (ZGA) in two phases: a minor wave in the 1-cell embryo and a major wave at the 2C stage ^22–24^. Zygotic factors necessary for the major wave of zygotic genome activation (ZGA) are expressed during the minor wave, such as *Dux* and the *OBOX* family genes, which exhibit partial redundancy ^25–29^.

The complexity of events during the few hours following fertilization and the limited amount of biological material make studying the molecular mechanisms linking genome organization to zygotic genome transcription extremely challenging. An *in vitro* alternative is the spontaneous conversion of a small fraction (<1%) of mouse embryonic stem cells (ESCs) into 2C-like cells (2CLCs), which recapitulate key features of the 2C embryo ^30^. These cells activate the transcription factor Dux, essential for triggering the expression of 2C-specific transcripts, such as the endogenous retrovirus *MERVL*, the *Zscan4a-f* gene family (referred to as Zscan4), *Tcstv1/3*, *Tdpoz1–5*, *Eif1a-like* cluster, *Nelfa*, and *Zfp352*^26,31–35^. Additionally, compared to ESCs, 2CLCs exhibit epigenetic characteristics typical of the 2C embryo, including higher histone mobility, increased chromatin accessibility, weaker genome-wide TAD organization, and reduced DNA methylation levels ^31,32,36–38^. Lastly, 2CLCs possess totipotent-like features, with a higher potential than ESCs to contribute to both embryonic and extra-embryonic lineages ^30^. However, significant differences exist between the *in vitro* model and *in vivo* 2C embryos, as 2CLCs do not fully replicate the natural embryo’s environment, progression, or gene expression patterns. Recently, specific treatments applied to ESCs induced 2CLCs that demonstrated a higher transcriptomic and epigenetic concordance with 2C embryos, reflecting a closer mimicry of the totipotent state ^39–42^.

This project aimed to use 2CLCs to conduct high-throughput genomic and epigenetic studies to better understand how 3D genome organization aligns with chromatin remodeling to enable the specific expression of ZGA genes while repressing others. In this work, we demonstrate that in spontaneous 2CLCs, the transient formation of Z-DNA (an alternative DNA conformation) is central for both establishing the totipotent-like transcriptional program and promoting cis- and trans-chromosomal interactions between genomic domains enriched in Zscan4-binding microsatellite repeats, defining a distinct nuclear compartment (Z compartment). Disrupting these specific interactions by depleting Zscan4 led to increased expression of genes involved in embryonic development and cell differentiation, indicating that the Z-compartment in 2CLCs has a repressive function that may be essential for safeguarding the totipotent-like state.

## Results

### 1. Identification of long-range chromatin interactions in 2CLCs

To explore the 3D genome architecture in 2CLCs, we utilized a previously published mouse ESC line with a td-Tomato transgene controlled by the MERVL promoter (MERVL::tdTomato), a reliable marker for the 2C-like state ^43^. The spontaneous Tomato-positive 2CLCs and Tomato-negative ESCs were purified by flow cytometry and subjected to Hi-C, a chromosome conformation capture-based method ^44^. Overall, chromosome organization remained largely similar between the two populations with prominent active/inactive (A-B) compartmentalization and TAD structures (**Fig. 1a, left**). Chromatin contact probabilities decreased similarly with increasing genomic distance, except at very long-range distances (>10 Mb), where a slight increase in contact frequencies was observed in 2CLCs compared to ESCs (**Extended Data Fig. 1a**). Deeper analysis of functional chromatin features revealed a weakening of TAD and loop strength in 2CLCs when averaged across all genomic positions (**Extended Data Fig. 1b, c**). Moreover, A-B compartmentalization remained largely similar between both cell types (**Extended Data Fig. 1d**), with a strong linear correlation between eigenvalues (r = 0.899). Consistent with previous reports ^31,32,38,45^, these data indicate that 2CLCs overall exhibit a more relaxed chromatin architecture than ESCs.

**Figure 1.**
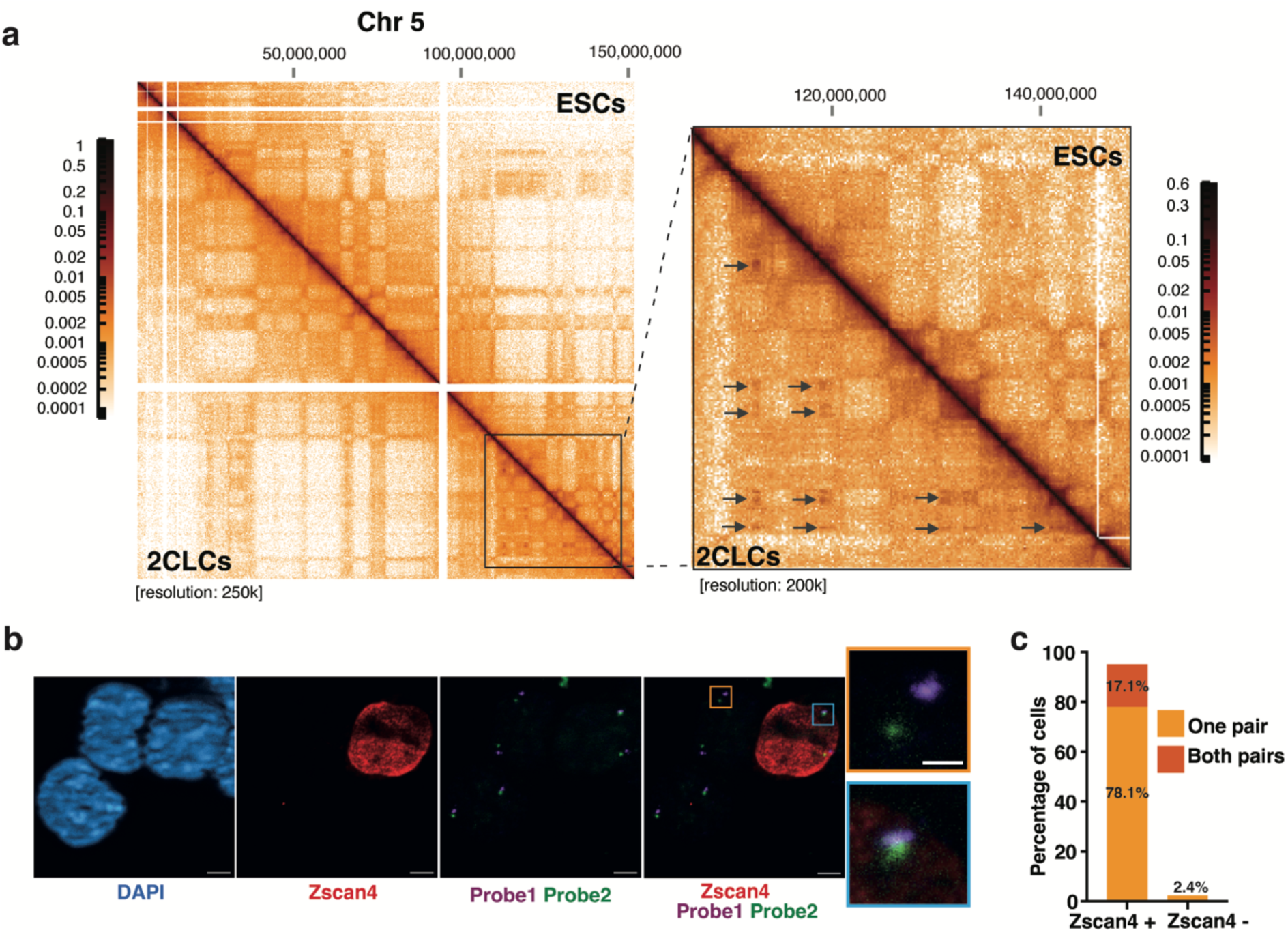
Identification of specific genomic interactions in 2CLCs. (**a**) Contact probability matrix for the entire chromosome 5 (Chr 5), comparing mouse ESCs (top right) and 2CLCs (bottom left), with a zoomed-in view (right) on one end of the chromosome. The arrows indicate 2CLC-specific chromatin interactions. (**b**) Representative images of DNA fluorescence *in situ* hybridization (DNA-FISH) combined with immunofluorescence in ESC cultures. 2CLCs were identified by Zscan4 staining, and DNA was counterstained with DAPI. Scale bar: 10 µm. The upper zoomed-in image shows non-overlapping probes in an ESC, the lower one shows overlapping probes in a Zscan4-positive 2CLC. Scale bar: 2 µm. (**c**) Quantification of immuno-DNA-FISH experiments showing the percentage of overlapping probes in Zscan4-positive 2CLCs (n = 41) and Zscan4-negative ESCs (n = 904) from four independent replicates.

Intriguingly, visual inspection of Hi-C maps revealed a distinct pattern in 2CLCs, absent in ESCs, characterized by extensive long-range interactions predominantly occurring at the distal ends of chromosomes such as 4, 5, 6, and 8, opposite to the centromere (**Fig. 1a, right, and Extended Data Fig. 1e**). Similar patterns were observed in 2CLCs when reanalyzing a published Hi-C dataset (**Extended Data Fig. 2a**)^38^. To investigate the spatial organization of these large interaction domains at the single-cell level, we performed DNA fluorescence in situ hybridization (FISH) with two DNA probes located 29 Mb apart on chromosome 5 (**Extended Data Fig. 2b**). The 2CLC population was detected in the ESC culture by immunolabeling the Zscan4 protein, a specific marker of 2CLCs ^30,34^. Strikingly, more than 95% of Zscan4-positive cells showed probe overlap, primarily at the nuclear periphery, on one or both copies of chromosome 5 (**Fig. 1b, c and Extended Data Fig. 2c)**. In contrast, only 2.4% of ECSs displayed a similar phenotype (**Fig. 1c**). We conclude that these interactions represent a novel and reproducible genomic feature of 2CLCs.

Next, to identify all the specific interacting regions genome-wide in 2CLCs, genomic TADs were first retrieved from the ESCs matrix using Chromosight ^46^, corresponding to 2,808 elements (**Supplementary Table 1**). For each intrachromosomal pair of TADs, a log2 ratio was calculated between counts for 2CLCs and ESCs. Applying a stringent cutoff (Log2 FC > 2.2), 106 TADs forming 85 specific TAD-TAD interactions (TAD-TADi) were identified, with many TADs interacting with multiple others (**Supplementary Table 2**). These TAD- TADi regions align well with the chromatin interactions visually observed in 2CLCs, notably at the end of chromosome 5 (**Extended Data Fig. 2d**), confirming the accuracy of the analysis.

Recent studies have shown that treating ESCs with specific compounds induces the emergence of 2CLCs with transcriptomic and epigenetic features more closely resembling 2C embryos compared to spontaneous 2CLCs ^39–41^. To test whether TAD-TADi are similarly observed in induced 2CLCs, MERVL::tdTomato ESCs were treated with Pladienolide B (PlaB), a spliceosome inhibitor, for 48 hours ^39^. As anticipated, PlaB treatment led to a roughly 15-fold increase in the emergence of 2CLCs (**Extended Data Fig. 2e, f**), and Hi-C experiments revealed that PlaB-induced 2CLCs exhibited TAD-TADi similar to those observed in spontaneous 2CLCs (**Extended Data Fig. 2g**), suggesting that these chromatin interactions may be a specific hallmark of totipotent-like cells.

Altogether, our data demonstrate local changes in chromosomal architecture during the transition from pluripotent to totipotent-like states, marked by the formation of long-range interactions in 2CLCs.

### 2. Formation of the Z-compartment in 2CLCs

To investigate the mechanisms driving the specific interactions identified in 2CLCs, we hypothesized that a chromatin factor uniquely expressed in 2CLCs could be the key regulator. We analyzed the contact probability between binding motifs of Dux and Zscan4, two main markers of 2CLCs ^26,34^. Interestingly, binding motifs for Zscan4, but not for Dux, showed a high frequency of contact probability in 2CLCs (**Fig. 2a, Extended Data Fig. 3a and Supplementary Table 3**). Therefore, we focused our analysis on the genomic loci bound by Zscan4 in 2CLCs ^34^, revealing that the distribution of Zscan4 peaks is biased toward one end of the chromosomes, opposite the centromere (**Fig. 2b and Extended Data Fig. 3b**), similar to the specific interactions observed in 2CLCs (**Fig. 1a and Extended Data Fig. 1e**). This non-random distribution is likely driven by the accumulation of its binding motif, corresponding to (CA/TG) dinucleotide repeats ^34^, at this chromosomal end (**Fig. 2c**). Analysis of genomic loci bound by Zscan4 in 2CLCs confirmed a specific enrichment of cis-contacts in 2CLCs compared to ESCs, along with unexpected trans-contacts between chromosomes (**Fig. 2d-f**). Given the annular staining of Zscan4 at the nuclear periphery in 2CLCs (**Extended Data Fig. 2c**), we hypothesized that the specific genomic interactions enriched for Zscan4 may be spatially segregated from the typical A-B compartmentalization in 2CLCs, occupying a distinct 3D nuclear space. For that, we employed a strategy similar to one previously reported for identifying a chromatin compartment induced by double-strand breaks ^47^. In brief, PCA was applied to the differential Hi-C map of the entire genome, comparing the tomato-positive 2CLC dataset to the tomato-negative ESC dataset, to extract the first eigenvector (PC1), which captures the largest variance (**Supplementary Table 4**). Genomic regions within the top 5% of positive PC1 values (corresponding to 617 bins of 200 kb) are strongly enriched for Zscan4 binding sites and were further referred to as the ’Z’ compartment (**Fig. 2f, g**). Indeed, Z compartment exhibited a 6.4-fold higher density of Zscan4 peaks (p-value < 1.e^-4^) than randomly shuffled peaks across the genome (**Extended Data Fig. 3c**). Moreover, we confirmed stronger intra- and inter-chromosomal interactions for the genomic regions associated with high PC1 values in 2CLCs compared to ESCs (**Extended Data Fig. 3d**). Of note, not all interacting TADs are part of the Z compartment (**Supplementary Tables 2, 4**), suggesting that a subset of TAD-TADi identified in 2CLCs may form independently of Zscan4-enriched regions. Interestingly, active hubs of inter-chromosomal interactions have been described in mouse ESCs using an alternative chromosome conformation capture method (SPRITE)^48^, which we found to largely overlap with the Z compartment displaying high PC1 values (**Extended Data Fig. 3e, f**). This suggests that the inter-chromosomal interactions exist in MERVL-negative ESCs (not detectable by us using Hi-C) and are strongly reinforced in MERVL-positive 2CLCs, or that these trans-contacts observed in the bulk ESC population by Quinodoz et al. ^48^ come exclusively from spontaneous 2CLCs.

**Figure 2.**
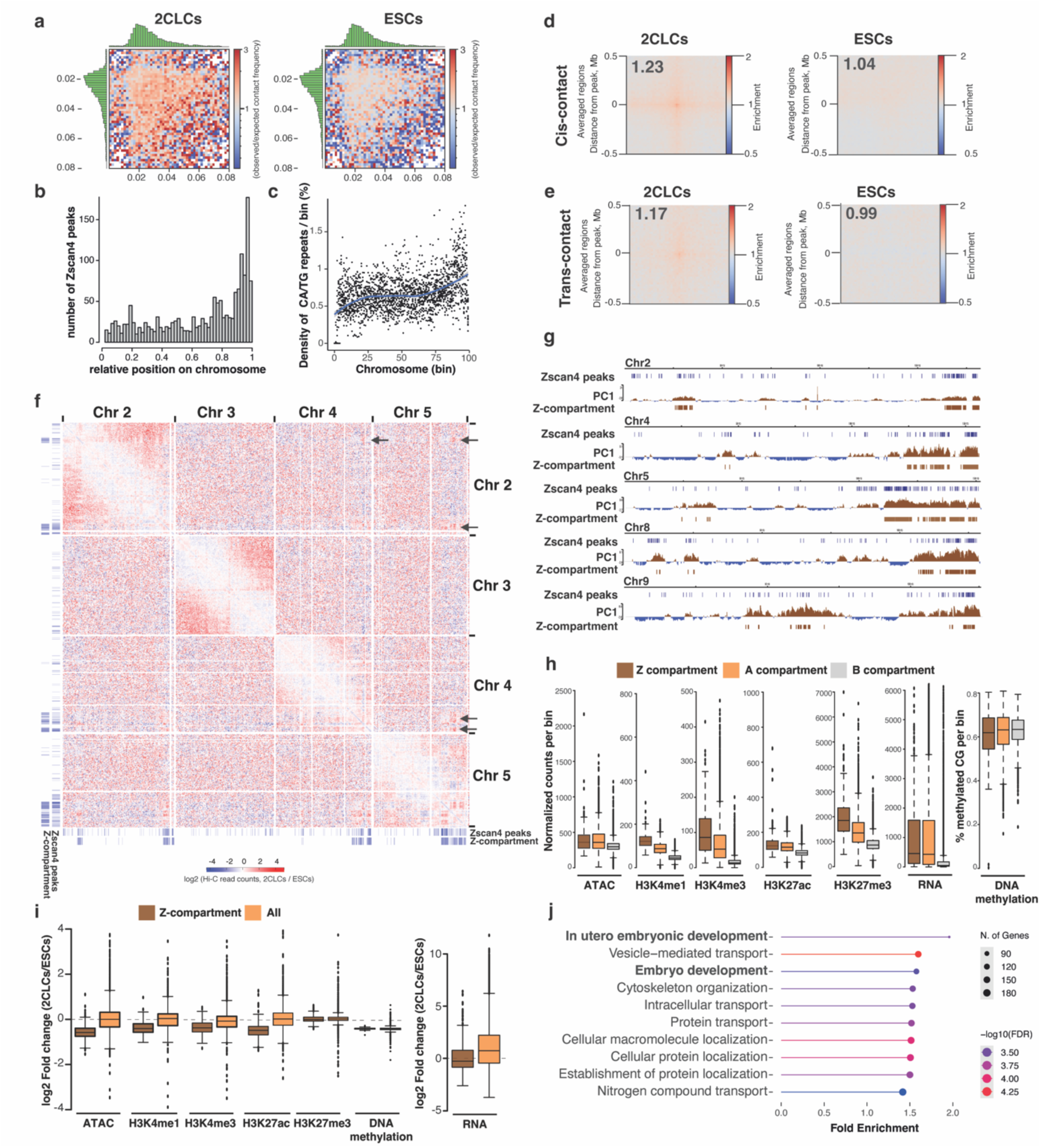
Identification and characterization of the Z-compartment in 2CLCs. (**a**) Saddle plot showing contact probability between 20kb genomic bins ranked according to the number of Zscan4 binding motifs per bin in 2CLCs (left) and ESCs (right). (**b**) Distribution of Zscan4 peaks along the normalized chromosome length. Each chromosome was divided into 50 bins of equal size. (**c**) Percentage of CA/TG repeats along the normalized chromosome length. Each chromosome was divided into 100 bins of equal size. For each bin, the percentage of CA/TG repeats is shown for the 19 mouse autosomes. (**d**) Pile up plot of cis-contact probability enrichment at Zscan4 binding sites in 2CLCs (left) and ESCs (right). (**e**) Same as ‘d’ for trans-contact. (**f**) Differential Hi-C contact matrix between 2CLCs and ESCs for chromosomes 2, 3, 4, and 5. Arrows indicate trans-contacts between Zscan4-enriched regions in the Z-compartment. The tracks for Zscan4 peaks and the Z-compartment are shown below and to the left of the matrix. Resolution: 1000kb. (**g**) Screenshots of Zscan4 peaks, PC1 from Hi-C analysis (2CLCs/ESCs), and the Z-compartment for chromosomes 2, 4, 5, 8, and 9. (**h**) Normalized counts per 200 kb bin for ATAC-seq, histone marks, and RNA-seq across the Z, A, and B compartments in 2CLCs. Percentage of methylated CG in the Z, A, and B compartments in 2CLCs. (**i**) Log2 fold change of ATAC-seq, histone marks, DNA methylation, and transcription signals per bin of 200kb between 2CLCs and ESCs in the Z-compartment and across the genome. (**j**) Ontology analysis of the biological processes associated with genes located in the z-compartment.

Characterization of the Z-compartment revealed that 93% of it (574 out of 617 bins of 200 kb) originates from compartment A in ESCs, while only 7% (43 out of 617) comes from compartment B. Next, to explore Z-compartment epigenetic and transcriptomic specificity, we compared the ATAC-seq, chromatin marks and RNA signals across A, B, and Z compartments in 2CLCs (**Fig. 2h and Supplementary Table 5**) ^38^. As expected, the A compartment exhibited higher levels of active marks, transcription, and chromatin accessibility than the B compartment. The Z compartment, enriched in Zscan4 peaks (**Extended Data Fig. 3g**), closely resembled the A compartment, showing similar ATAC-seq signal, H3K27ac levels, and transcription activity. However, it displayed a notable increase in H3K4me3, H3K4me1, as well as the repressive mark H3K27me3. DNA methylation levels remained largely unchanged across compartments. Finally, investigating the dynamics of these marks during the pluripotent-to-totipotent transition revealed a significant decrease in the Log2 fold change of both chromatin accessibility and the active histone marks H3K4me1, H3K4me3, and H3K27ac in 2CLCs compared to ESCs, with no changes observed for the repressive mark H3K27me3 (**Fig. 2i**). The dynamic changes observed in the Z compartment were not detected genome-wide (All) (**Fig. 2i**). Regarding DNA methylation levels, we confirmed the hypomethylated status of 2CLCs compared to ESCs ^31^, but no differences were detected between the Z compartment and the rest of the genome (**Fig. 2h**). Finally, transcription levels in the Z compartment were reduced in 2CLCs compared to ESCs, and ontology analysis revealed that genes within the Z compartment were enriched for biological processes associated with embryonic development (**Fig. 2i, j**).

These data indicate that genomic interactions within the Z compartment in 2CLCs are linked to localized epigenomic modifications, leading to less active and less transcribed regions in 2CLCs than in ESCs.

### 3. The chromatin factor Zscan4 is essential for the interactions specific to 2CLCs

To determine whether Zscan4 is required for the formation or maintenance of the specific interactions enriched in the Z compartment in 2CLCs, Zscan4 was depleted by up to 87% using siRNAs targeting the different isoforms (**Fig. 3a and Extended Data Fig. 4a, b**). Importantly, Zscan4 knockdown neither affected the percentage of MERVL::tdTomato-positive cells as detected by flow cytometry nor significantly altered the 2C transcriptional program compared to control 2CLCs (except for the Zscan4 family) (**Extended Data Fig. 4b, c**), consistent with a previous report ^45^. Analysis of Hi-C data in 2CLCs revealed that the strong Zscan4 reduction abolished cis- and trans-interactions between Zscan4 binding sites across the genome, unlike the control condition (**Fig. 3b, c and Extended Data Fig. 4d**). To investigate the functional role of Zscan4-dependent interactions, transcriptomic analysis was performed in Zscan4-depleted 2CLCs. Zscan4 suppression upregulated 1,857 transcripts, which were not randomly distributed across the genome but instead formed 24 clusters (**Extended Data Fig. 4e and Supplementary Table 6**). These clusters were highly enriched in Zscan4 peaks, notably at chromosome ends (e.g., chromosomes 4, 5, and 11) (**Fig. 3d and Extended Data Fig. 4e)**. In contrast, 1,464 downregulated genes formed only 4 clusters lacking Zscan4 peak enrichment (**Extended Data Fig. 4e, f and Supplementary Table 6**). Ontology analysis revealed that upregulated genes, but not the downregulated, were associated with development-related terms such as “morphogenesis of an epithelium”, “tube development”, “embryo development”, “neurogenesis” (**Fig. 3e and Extended Data Fig. 4g**). These data suggest that in absence of Zscan4, 2CLCs overexpressed genes associated with later stages of development, beyond the 2C stage. To test this hypothesis, we used a recent transcriptomic dataset tracking nascent transcript dynamics from the zygote to the blastocyst stage ^49^. Clustering analysis by K-means identified six distinct clusters grouping the expression of 18,508 genes, each reflecting unique developmental expression dynamics (**Extended Data Fig. 4h, i**). As postulated, the upregulated genes in Zscan4-depleted 2CLCs exhibited a distinct distribution compared to the 18,508 genes, with a significant enrichment in the later stages of pre-implantation development (4C, 8C, and morula/blastocyst stages) (**Fig. 3f**).

**Figure 3.**
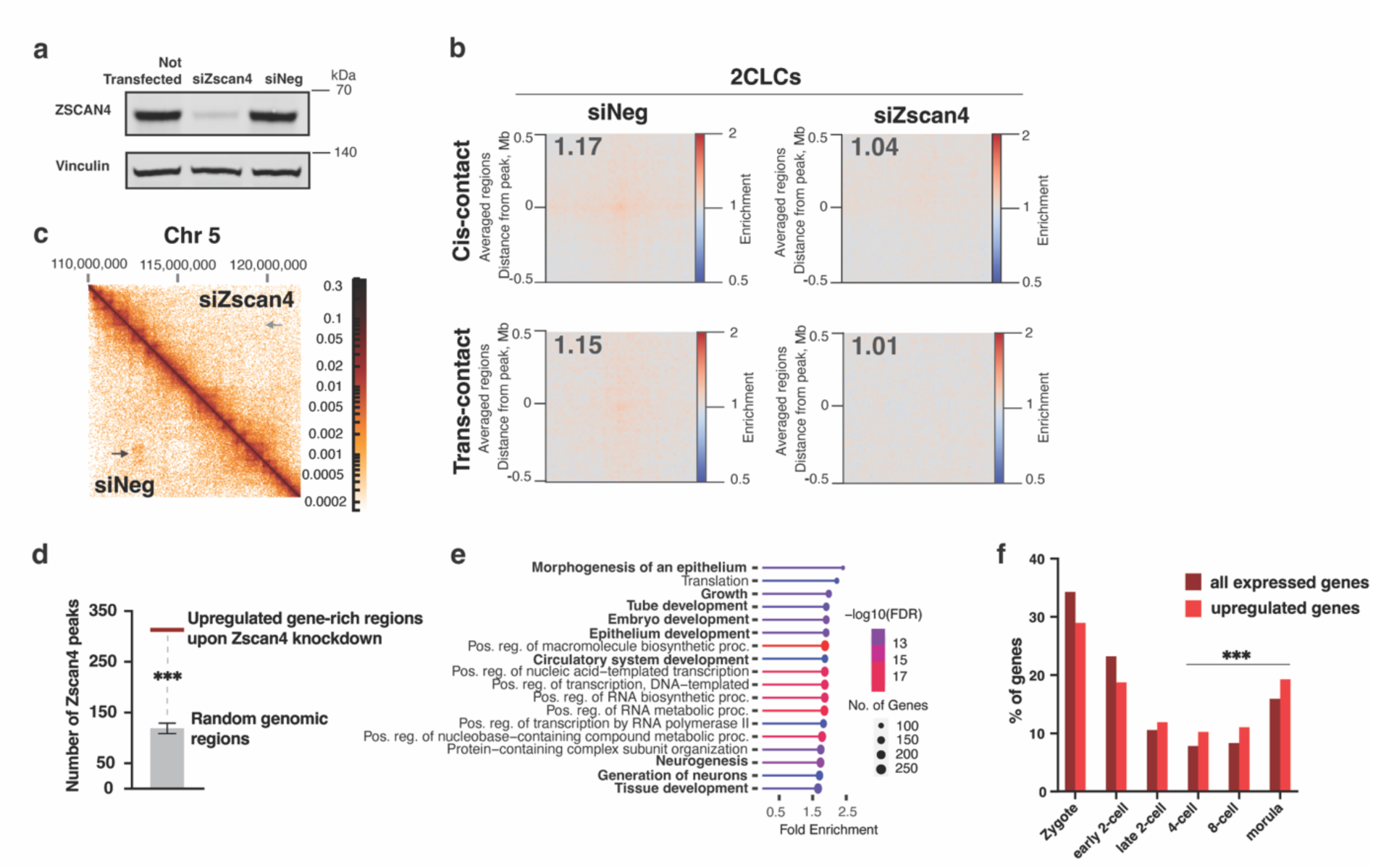
Zscan4 is required for the specific genomic interactions characteristic of 2CLCs. (**a**) Representative example of immunoblots in ESCs for Zscan4 and Vinculin, used as the housekeeping control; siZscan4: siRNAs against Zscan4, siNeg: non-targeting siRNA control. (**b**) Pile up plot of cis- and trans-contact probability enrichment at Zscan4 binding sites in Zscan4-depleted 2CLCs (right) or in control 2CLCs (siNeg, left). (**c**) Hi-C map showing a 2CLC-specific chromatin interaction on chromosome 5 in control 2CLCs (black arrow) and its absence in Zscan4-depleted 2CLCs (gray arrow). (**d**) Number of Zscan4 peaks in upregulated gene-rich regions upon Zscan4 knockdown in 2CLCs (red bar) in comparison to random regions (1000 sets of random regions were tested, p < 1e^-4^). (**e**) Ontology analysis of the biological processes associated with upregulated genes upon Zscan4 knockdown in 2CLCs compared to control 2CLCs. (**f**) Distribution of upregulated genes upon Zscan4 knockdown in 2CLCs compared to the distribution of all genes expressed during early development, from the zygote to the morula stage. Each gene was assigned to a specific developmental stage using the k-means method. A Chi-square test was performed on the number of upregulated genes at the 4-cell, 8-cell, and morula stages compared to the total number of expressed genes at these stages (p = 9.91e-7).

Altogether, we propose an unanticipated role for Zscan4 in shaping 3D chromatin organization in totipotent-like cells by promoting compartmentalization of Zscan4-dependent chromatin interactions in 2CLCs, which act as transcriptional repressors of development-related genes.

### 4. Z-DNA formation induces totipotent-like cells

Zscan4 has been shown to bind dinucleotide (CA)n tandem repeats to prevent the formation of Z-DNA, a left-handed helical conformation of DNA associated with genomic instability ^34^. Consistent with this, we observed an important overlap between Z-DNA motifs ^50^ and Zscan4 peaks in 2CLCs, with a linear correlation of r = 0.36 between the two genomic tracks (**Extended Data Fig. 5a**). Similar to Zscan4 motifs, 2CLCs exhibited an increased contact probability between Z-DNA-prone regions compared to ESCs (**Extended Data Fig. 5b and Fig. 2a**). Building on previous studies showing that Z-DNA induction activates p53 in cancer cell lines ^51,52^ and findings from mouse ESCs indicating that p53 induces Dux, the master regulator of the totipotency-associated program ^53^, we hypothesized that Z-DNA formation might be crucial for the emergence of 2CLCs. Using a validated anti-Z-DNA antibody ^54^ alongside an anti-Nelfa antibody as one of the earliest 2CLC marker ^33^, we observed diffuse nuclear Z-DNA staining in 40.2% and 32.4% of Nelfa- positive 2CLCs from two different ESC lines, a feature absent in Nelfa-negative ESCs (**Fig. 4a**). Furthermore, most Z-DNA-positive cells lacked Zscan4 expression, a more downstream 2CLC marker, with only a 4% overlap in the E14-ESC line (4 Z-DNA-positive cells out of 101 counted Zscan4-positive cells, N=3) (**Fig. 4b**). Of note, the few Z-DNA- and Zscan4-double-positive cells detected were adjacent to one another, suggesting that this intermediate state happens shortly after mitosis. Altogether, these data suggest that formation of Z-DNA is a transient feature of early 2CLCs.

**Figure 4.**
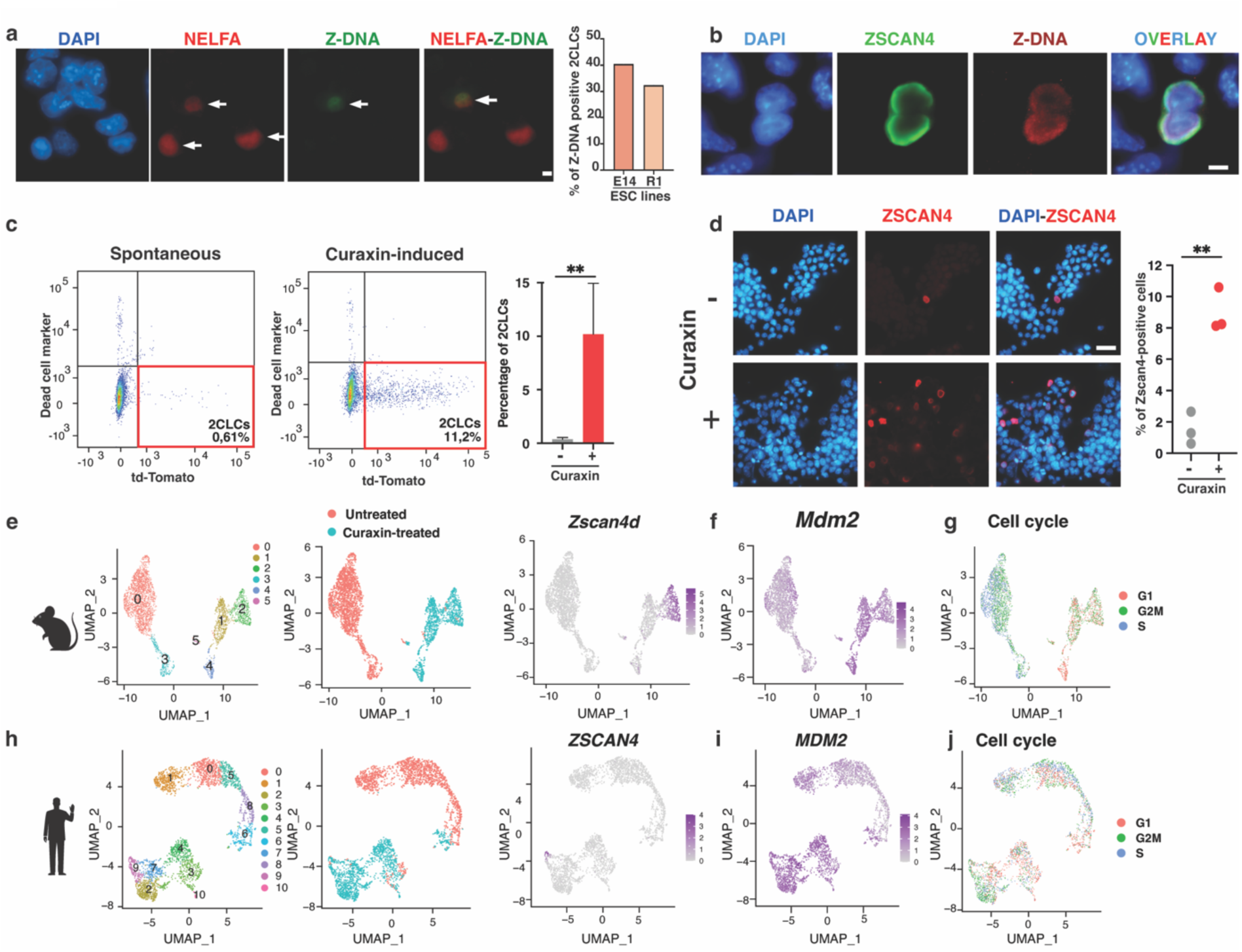
Curaxin-induced Z-DNA formation increases both the emergence of murine 2CLCs and human 8CLCs. (**a**) Representative immunofluorescence images showing 2CLCs detected by an anti-NELFA antibody (red) and Z-DNA stained with an anti-Z-DNA antibody (green). DNA is counterstained with DAPI. The graph displays the percentage of 2CLCs positively stained for Z-DNA in two murine ESC lines: n=52 2CLCs from E14 ESCs, n=50 2CLCs from R1 ESCs, N=2 independent experiments. Scale bar: 5 μm. (**b**) Representative immunofluorescence images showing 2CLCs detected by an anti-Zscan4 antibody (green) and Z-DNA stained with an anti-Z-DNA antibody (red). DNA is counterstained with DAPI. n=50 2CLCs from E14 ESCs, N=2 independent experiments. Scale bar: 5 μm. (**c**) Representative dot plots from flow cytometry analysis showing an increase in the MERVL-tdTomato-positive 2CLC population (red square) after 48 hours of treatment with 0.5 μM curaxin, compared to untreated mESCs. The graph depicts the quantification from N=6 independent experiments, ** p<0.01, two-tailed t-test. (**d**) Representative immunofluorescence images showing an increase in the number of 2CLCs stained with an anti-Zscan4 antibody (red) in mESCs treated with 0.5 μM curaxin for 48 hours, compared to the control condition, and the associated quantification for N=3 independent experiments, ** p<0.01, two-tailed t-test. Scale bar: 50 μm. (**e**) Uniform Manifold Approximation and Projection (UMAP) plots of 3,752 murine ESCs treated or untreated with 0.5 μM Curaxin for 48 hours. Cells are colored by their cluster annotation (left), treatment condition (middle), and *Zscan4d* expression (right). (**f**) UMAP plot as in ‘e’ showing the expression of *Mdm2*. (**g**) UMAP plot as in ‘e’ showing cell cycle phases. (**h**) as in ‘e’ for 3,131 human induced pluripotent cells treated or untreated with 0.5 μM curaxin for 48 hours. (**i**) UMAP plot as in ‘h’ showing the expression of *MDM2*. (**j**) UMAP plot as in ‘h’ showing cell cycle phases.

To explore the potential role of Z-DNA in initiating the pluripotent to totipotent-like transition, ESCs were treated with curaxin (0.5μM) for 2 days, a Z-DNA-inducing compound ^55,56^. Remarkably, curaxin treatment of ESCs for 48 hours led to a massive increase in MERVL::tdTomato-positive cells, with their proportion rising from 0.36% in untreated conditions to 10.18%, as measured by flow cytometry (**Fig. 4c**). Immunostaining experiments confirmed the strong increase in 2C markers such as Zscan4 and Nelfa following curaxin treatment (**Fig. 4d and Extended Data Fig. 5c**). Next, single cell transcriptomic experiments (scRNA-seq) were performed in curaxin-treated and control condition in ESCs. Clustering analysis revealed a total of six different cell populations, with cluster 2 showing specific expression of the top 100 markers of the 2CLC signature, such as the *Zscan4*, *Usp17l*, *Tdpoz*, and *Pramel* families (**Fig. 4e, Extended Data Fig. 5d, e and Supplementary Table 7**). Strikingly, cluster 2 was almost exclusively composed of curaxin-treated cells (474 out of 483), with curaxin treatment increasing the proportion of 2CLCs by 96-fold, accounting for 35.7% of the treated ESC population (**Fig. 4e**). Moreover, the treatment drove a global transcriptional shift, with the emergence of clusters 1 and 4 containing almost exclusively curaxin-treated cells (**Fig. 4e**). Ontology analysis of the markers of clusters 1 and 4 revealed significant enrichment of terms related to p53 signaling, cell cycle, and DNA replication (**Extended Data Fig. 5f**). The increased expression of the two main p53 targets, *Cdkn1a* and *Mdm2*, as well as the sharp increase in p53 protein and phospho-p53 in curaxin-treated ESCs (**Fig. 4f, Extended Data Fig. 5g, h**), confirmed that curaxin-induced Z-DNA activates the p53 pathway. To analyze the impact of curaxin treatment on the cell cycle, a cell cycle phase score was assigned to each cell from the scRNAseq dataset based on canonical G1, S, G2/M markers. Control ESCs displayed a typical cell cycle distribution characterized by a short G1 phase, whereas curaxin-treated cells showed a marked decrease in S-phase cells (12-fold reduction) and an accumulation in G1 phase relative to controls (**Fig. 4g**). This observation is interesting for two reasons: first, reduced DNA replication fork speed has been shown, both *in vitro* and *in vivo*, to promote the emergence of totipotent and totipotent-like cells ^57^; second, DNA replication initiation zones identified in ESCs prominently feature a motif composed of successive (CA) microsatellite repeats ^58^. This suggests that Z-DNA formation might influence DNA replication timing. Taken together, these findings indicate that Z-DNA induction by curaxin led to the emergence of 2CLCs with a transcriptomic signature similar to that of spontaneous 2CLCs. Moreover, Hi-C experiments in curaxin-induced 2CLCs revealed comparable cis- and trans-interactions around Zscan4 peaks (**Extended Data Fig. 5i**), demonstrating that their 3D genome organization is also consistent with that of spontaneous 2CLCs.

Recently, a novel *in vitro* model called 8-cell-like cells (8CLCs) was identified, which mimics the transcriptional features of human 8-cell embryos, a stage analogous to the mouse 2C stage and marked by ZGA ^59^. These cells express key ZGA markers such as *ZSCAN4*, *LEUTX*, *KLF17*, and *TPRX1* ^59^. Leveraging this model, we treated human induced pluripotent stem cells (iPSCs) with curaxin to investigate whether Z-DNA induction also contributes to the emergence of totipotent-like cells. The 8CLC population was clearly identified in cluster 9 by the expression of the top 100 human-specific totipotent markers ^59^, including *ZSCAN4*, *KLF17*, and *TPRX1* (**Fig. 4h, Extended Data Fig. 5j, k and Supplementary Table 8**). Cluster 9 comprised only 6 cells out of 1,664 from the control condition, compared to 148 cells out of 1,467 from the curaxin-treated condition, highlighting a massive increase in the proportion of totipotent-like cells following treatment (**Fig. 4h**). As observed in mice, curaxin treatment increased the expression of genes involved in p53 signaling, including the overexpression of *CDKN1A* and *MDM2*, and reduced the proportion of curaxin-treated iPSCs in the S phase from 36.8% in control iPSCs to 18.3% (**Fig. 4i, j and Extended Data Fig. 5l, m**). We conclude that the formation of Z-DNA may represent a conserved mechanism for initiating the pluripotent-to-totipotent conversion in both murine and human cells.

Altogether, these data support a model in which the formation of Z-DNA acts as an upstream event that activates p53, leading to the induction of Dux and the subsequent establishment of the totipotent-like transcriptional program.

### 5. Inhibiting the polyamine pathway prevents Z-DNA formation and disrupts Zscan4-dependent compartmentalization in 2CLCs

We have demonstrated that the formation of Z-DNA in 2CLCs foster the emergence of totipotent-like cells. Furthermore, Zscan4 binds to chromatin at motifs prone to forming Z-DNA ^34^, promoting interactions between Zscan4-rich domains that repress gene expression in 2CLCs. We now aim to establish a direct link between Z-DNA and the specific interactions mediated by Zscan4 in 2CLCs. Various molecular and environmental factors, including high polyamine concentrations, have been described to favor transitions to Z-DNA ^60^. Natural polyamines, such as putrescine, spermine, and spermidine, are polycationic molecules that interact with negatively charged DNA to induce conformational transitions and stabilize Z-DNA. Interestingly, a metabolomic study revealed elevated polyamine levels in 2C embryos and 2CLCs compared to blastocysts and ESCs, respectively ^61^. To test whether reducing polyamine levels prevents Z-DNA formation in 2CLCs, mouse ESCs were treated with a combination of difluoromethylornithine (DFMO), an irreversible polyamine synthesis inhibitor, and AMXT-1501 tetrahydrochloride, a potent polyamine transport inhibitor (**Extended Data Fig. 6a**) ^62,63^. As anticipated, polyamine concentrations dramatically dropped in ESCs 48 hours post-treatment (**Extended Data Fig. 6b**). This reduction was accompanied by the complete loss of Z-DNA staining in Nelfa-positive 2CLCs and a 46% decrease in the percentage of spontaneous 2CLCs (**Fig. 5a, b and Extended Data Fig. 6c**), demonstrating that polyamine inhibition effectively prevents Z-DNA formation and alters cell fate conversion to totipotent-like cells. Remarkably, Hi-C experiments conducted on emerging 2CLCs at 24h and 48h under polyamine inhibition revealed a gradual dissolution of Zscan4-dependent chromatin interactions across the genome, as exemplified at the end of chromosome 5 (**Fig. 5c and d)**. Finally, we analyzed transcriptional variations following 48 hours of polyamine inhibition and identified 2,175 upregulated genes enriched for biological processes related to protein transport and catabolism, while 2,302 downregulated genes were associated with ribosome biogenesis in 2CLCs (**Extended Data Fig. 6d and Supplementary Table 6**). The downregulated genes clustered in 22 genomic regions enriched for Zscan4 binding, 12 of which overlapped with clusters containing upregulated genes in Zscan4-depleted 2CLCs (**Extended Data Fig. 6e, f**). These surprising findings suggest that disrupting totipotent-associated chromatin interactions by blocking Z-DNA formation led to transcriptional repression, whereas Zscan4 depletion increased transcription in these regions. Of note, upregulated genes formed only two significant clusters depleted of Zscan4 (**Extended Data Fig. 6e, f**).

**Figure 5.**
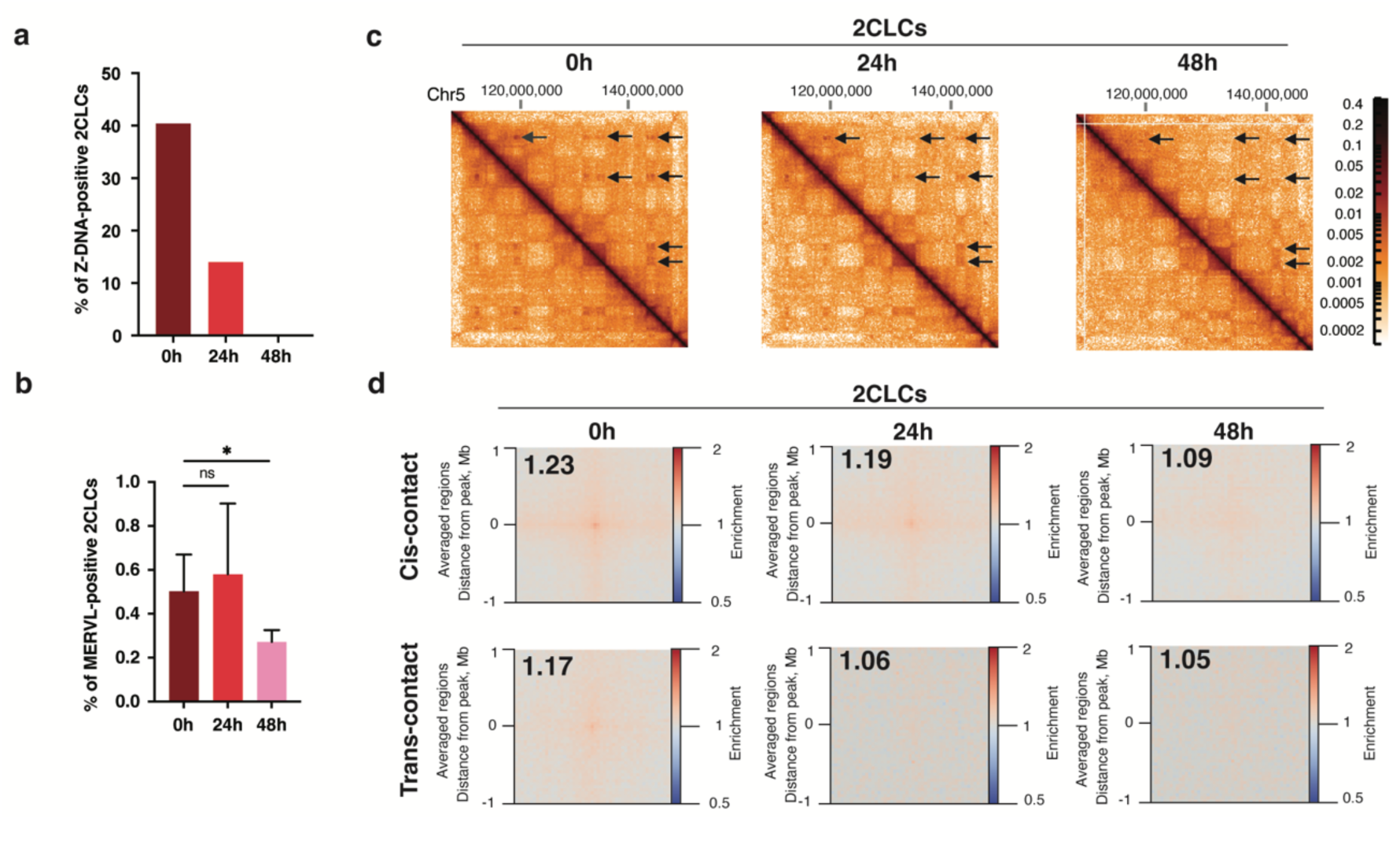
Polyamine synthesis inhibition disrupts Zscan4-dependent compartmentalization in 2CLCs. (**a**) Bar graph showing the percentage of Nelfa-positive 2CLCs displaying Z-DNA, determined by immunostaining experiments as in Figure 3a, at 0h, 24h, and 48h post-polyamine inhibition (0.25 mM DFMO and 15 μM AMXT-1501). Analysis of n=52 2CLCs at 0h, 53 at 24h and 59 at 48h from N=3 independent experiments. (**b**) Bar graph showing the percentage of MERVL-positive 2CLCs observed post-polyamine inhibition (0h, 24h, and 48h), as determined by flow cytometry. N=5 independent experiments; ns, not significant; * p < 0.05, two-tailed t-test. (**c**) Hi-C contact probability matrix for the indicated portion of the chromosome 5 (Chr 5), comparing spontaneous 2CLCs and 2CLCs 24h and 48h post polyamine inhibition (0.25 mM DFMO and 15 μM AMXT-1501). Arrows point to 2CLC-specific chromatin interactions identified in spontaneous 2CLCs. (**d**) Pile up plots of cis- and trans-contact probability enrichment at Zscan4 binding sites in spontaneous 2CLCs and 2CLCs 24h and 48h post polyamine inhibition (0.25 mM DFMO and 15 μM AMXT-1501).

In summary, we propose that elevated natural polyamines in 2CLCs may trigger the formation of Z-DNA, which is essential for promoting Zscan4-mediated interactions and regulating the expression of genes within these domains.

### 6. Chromatin and transcriptomic dynamics in the Z-compartment

To summarize the connections between the genomic elements associated with the Z compartment and the global epigenetic and transcriptomic dynamics during the pluripotent-to-totipotent-like transition, a hierarchical correlation matrix was generated genome-wide (**Supplementary Table 5**). The analysis revealed a positive correlation between the PC1 from Hi-C (2CLCs vs. ESCs), which defines the Z compartment (**Fig. 2**), and the enrichment of Zscan4 peaks ^34^, Z-DNA motifs, and DNA replication initiation zones ^58^, all of which share similar (CA) repeat motifs (**Fig. 6a, b and Extended Data Fig. 7a, b**). Of note, DNA methylation dynamics between 2CLCs and ESCs were tightly correlated with these elements. In contrast, genomic features of the Z compartment were negatively correlated with changes in DNA accessibility (ATAC), active histone marks, and transcription between 2CLCs and ESCs, while H3K27me3 dynamics appeared independent ^38^, suggesting that this repressive histone mark does not play a major role in Z compartment regulation.

**Figure 6.**
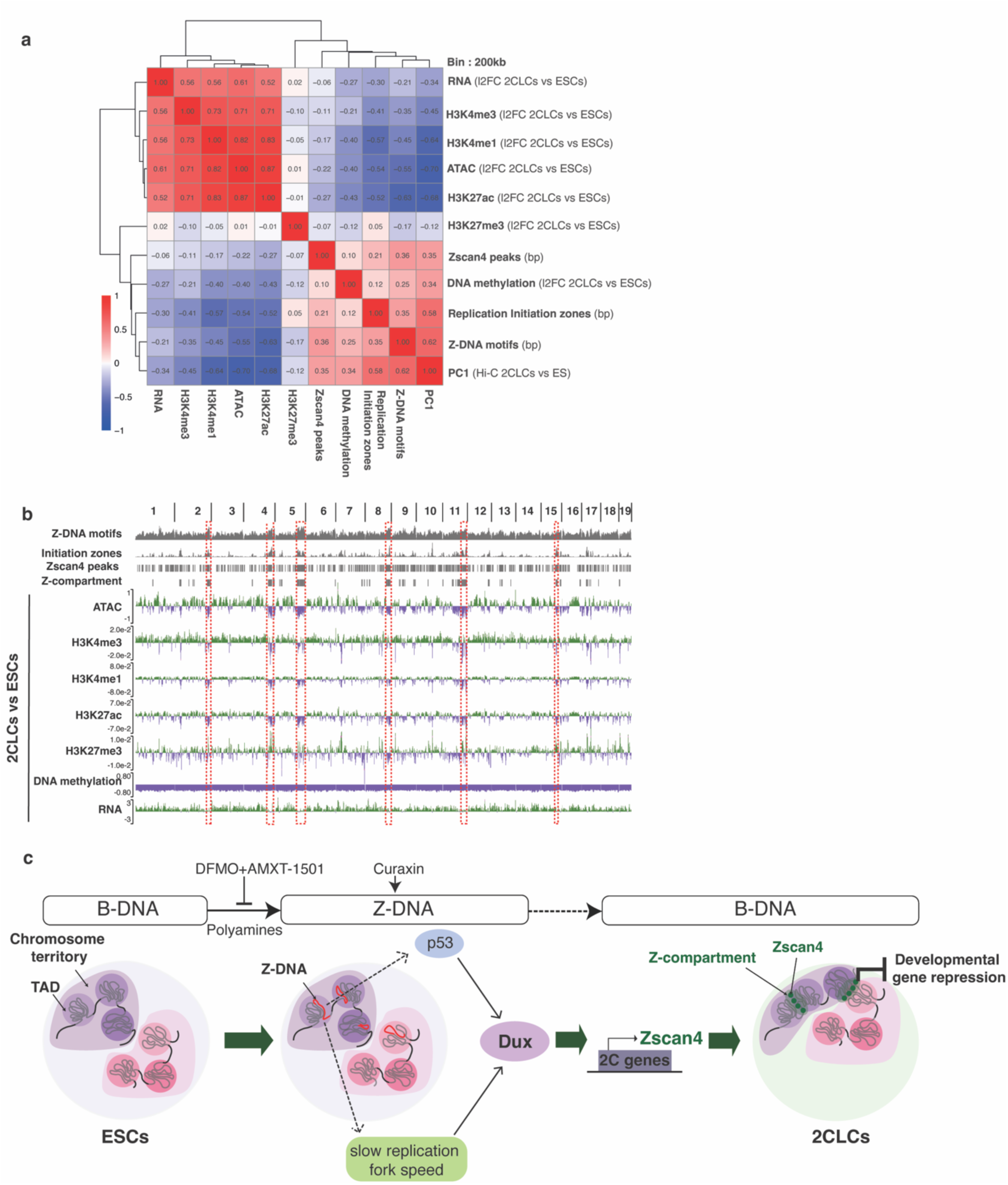
Epigenetic and transcriptomic dynamics during the pluripotent-to-totipotent-like conversion. (**a**) Hierarchical Spearman correlation matrix of the indicated elements. For histone marks, DNA methylation, and RNA transcription, the log2 fold change of the signal was calculated between 2CLCs and ESCs for each 200 kb bin. The eigenvector value (PC1) per 200 kb bin, derived from Hi-C analysis comparing 2CLCs to ESCs (as described in Figure 2), was used. Additionally, the percentage of bases corresponding to Z-DNA motifs, Zscan4 peaks, or DNA initiation zones was calculated per 200 kb bin. (**b**) Screenshot of Z-DNA motifs, DNA replication initiation zones, Zscan4 peaks, Z-compartment, and the log2 fold change of histone marks, DNA methylation, and RNA transcription signals between 2CLCs and ESCs across the 19 autosomal chromosomes. (**c**) Model illustrating the role of Z-DNA in the pluripotent-to-totipotent-like transition and its contribution to the compartmentalization of Zscan4-dependent interactions specific to 2CLCs. Mechanisms to be explored further are represented by dashed arrows. The scheme is simplified for clarity, and A-B compartmentalization is not shown.

These findings highlight the coordinated interplay between genome architecture, chromatin regulation, and transcription in defining the Z compartment during the totipotent-like transition.

## Discussion

In this study, we propose that an elevation in polyamines triggers Z-DNA conversion, which acts as a catalyst to induce *Dux* and establish the 2C transcriptional program, and as a seed to cluster Zscan4-mediated chromatin interactions, restricting the expression of development-associated genes (**Fig. 6c**).

Two primary mechanisms induce *Dux* expression: the activation of p53 and the slowing of DNA replication fork speed ^53,57^. Our findings, along with evidence from other studies in diverse cellular models, demonstrate that curaxin-induced Z-DNA formation increases both p53 levels and its active phosphorylated form ^51,52^. While Z-DNA formation does not seem to directly cause DNA damage ^51^, it alters the DNA helix by increasing the inter-base-pair distance and rearranging the sugar-phosphate backbone into a zigzag pattern ^64^. This structural change results in the unwrapping of DNA from histone octamers and destabilizes nucleosomes, creating topological stress able to activate p53 ^51,55^. An alternative pathway for p53 activation may involve the recruitment of the DNA repair machinery. For example, the mismatch repair complex Msh2-Msh3 has been shown to recognize Z-DNA and to recruit the nucleotide excision repair complex ERCC1-XPF, which cleaves Z-DNA, leading to genomic instability ^65^. This suggests that Z-DNA formation may indirectly induce DNA strand breaks, thereby activating ATM/ATR kinase-dependent phosphorylation of p53. Furthermore, Z-DNA directly influences DNA replication, as a recent study showed that Z-DNA/Z-RNA hybrids block replication ^66^, supporting our findings that curaxin treatment in ESCs disrupts entry into the S-phase. The formation of Z-DNA may thus activate *Dux* expression through both the activation of p53 and the regulation of DNA replication fork speed (**Fig. 6c**).

Z-DNA-positive 2CLCs represent an early transient state, suggesting that additional chromatin factors are required to convert Z-DNA back to B-DNA and to recruit Zscan4, as Zscan4 binds only to B-DNA, not Z-DNA ^34^. Several proteins, most of them containing a z-alpha domain, have been shown to specifically bind to Z-DNA such as Zbp1 and Adar1 but only Zbtb43 has been proven to bind and eliminate Z-DNA structure in germ cells of the mouse male fetus ^56,67–69^. Further studies are needed to understand the Z-DNA dynamics during the pluripotency to totipotency-like reversion.

Zscan4 is essential for maintaining genome stability, telomere elongation, and proper embryonic development in mice ^34,70,71^. Zscan4 is expressed during the late 2C embryo stage, and its reduction delays the transition to the 4-cell stage and lowers the success of implantation into the uterine epithelium ^70^. The totipotent-like cells are not identical to 2C embryos undergoing the natural ZGA process. Nevertheless, our data showing that Zscan4 is essential for repressing differentiation genes through the formation of a distinct chromatin compartment (Z compartment) involving intra- and inter-chromosomal interactions may represent a potential mechanism that could explain the defects in early developmental progression and implantation of embryos with decreased Zscan4 levels. Interestingly, recent findings indicate that Zscan4 associates with repressive chromatin linked to KAP1, Setdb1, LSD1, and HDAC1, promoting its disassembly and thereby increasing chromatin accessibility, which facilitates the transition to the 2CLC state ^72^. This shows that, depending on the epigenetic context, Zscan4 functions as a chromatin remodeler that can either open chromatin, as in the example mentioned above, or modify chromosome architecture to confine regions within a specific nuclear compartment to regulate gene transcription. Beyond its role as a chromatin dynamics modulator, Zscan4 binding may prevent the re-formation of Z-DNA, which could prolong the 2C state or even play a role in Z-DNA repair. Indeed, Zscan4 has been shown to interact with the repair factor PARP1, facilitating DNA repair in mouse ESCs ^73^. The notion that Zscan4 is associated with the DNA repair machinery is compelling, as the induction of double-strand breaks drives the compartmentalization of damaged regions away from conventional A-B domains ^47^.

The specific interactions dependent on Zscan4 we detected in 2CLCc are not randomly distributed along the chromosome. Zscan4 binds to the TGCACAC motif ^34^ primarily at the centromere-distal end regions, driven by the uneven localization of CA/TG repeats along the chromosomes. This indicates that specific genomic sequences bound by chromatin factors may co-segregate from the typical A-B compartmentalization to form higher-order inter-chromosomal contacts critical for gene regulation. In mouse ESCs, two hubs of inter-chromosomal interactions have been identified: one in proximity to the nucleolus and the other around nuclear speckles, associated with inactive and active transcription, respectively ^48^. Intriguingly, a large genomic fraction of the active hub found in ESCs overlaps with the Zscan4-dependent chromatin interactions identified in 2CLCs, forming the repressive Z-compartment. Consistently, these regions, which exhibit an open chromatin state, high levels of active histone marks, and strong transcriptional activity in ESCs, become less accessible in 2CLCs, losing active histone marks and displaying reduced transcriptional activity. A detailed spatiotemporal analysis by live imaging or DNA-FISH of these regions across cell states could provide deeper insights into the dynamic aspects of gene regulation in the nucleus. In this study, we found that the depletion of Zscan4 in 2CLCs increases the expression of gene clusters enriched in the Z-compartment. Conversely, preventing Z-DNA formation, an upstream event that similarly disrupts Zscan4-dependent interactions, decreases gene expression compared to control 2CLCs. This suggests that, in the absence of Z-DNA, a compensatory mechanism -likely epigenetic-occurs during the conversion of 2CLCs, strongly suppressing gene expression.

Numerous small molecule compounds targeting specific signaling pathways, epigenetic modifiers, or metabolic processes have been shown to massively induce the conversion to 2CLCs ^39–42,74,75^. In this study, we focused on investigating the spontaneous transition from ESCs to 2CLCs to avoid perturbing the system with non-specific drug effects that could alter chromosomal conformation, the epigenome, or the transcriptome. Therefore, the Z-DNA/Z compartment axis during the ESC-to-2CLC transition represents a dynamic process of chromosomal self-organization. Extrapolating this finding to an *in vivo* context during early development will require genome-wide high-throughput interaction data and local DNA-FISH to visualize long-range 3D structures in early embryos of both mice and humans. Indeed, we showed that curaxin-induced Z-DNA strongly increases both the proportion of 2CLCs and 8CLCs. The nuclear architecture of the early embryo is characterized by a dynamic post-fertilization reorganization of the chromatin conformations with a progressive establishment of compartmentalization. Although totipotent cells exhibit a permissive chromatin state and lack TADs, early repressive compartments such as LADs, detected as early as the zygote, and PADs, observed at the 2C stage, function as transcriptionally repressive genomic hubs ^12–14,20,21^. Similarly, the potential formation of the Z-compartment in totipotent cells may represent a new regulatory layer that prevents the expression of non-ZGA genes by spatially constraining specific chromosomal regions enriched in microsatellite repeats.

## Materials and Methods

### Cell culture and treatments

ES-E14 MERVLtd-tomato ES cell line, kindly gifted by Wolf Reik, Babraham Institute, Cambridge, UK were used for most of the experiments ^43^. Cells were maintained in Serum+Lif (S/L) medium (KnockOut DMEM supplemented with 15% ES cell qualified FBS, 1% GlutaMAX, 1% MEM non-essential amino acids, 1% penicillin-streptomycin, 0.1 mM 2-mercaptoethanol, 10 ng/mL Lif (Miltenyi Biotec, Cat #130-099-895) on gelatine-coated plates in a humidified incubator (37°C, 5% CO2). ES R1 cell line was maintained under the same conditions. Human iPS cell line was generously gifted by Prof. Dr. Nikolaus Rajewsky, Max Delbruck Center for Molecular Medicine, Berlin, Germany and corresponded to Gibco™ Human Episomal iPSC Line TMOi001-A (Cat# A18945). HuIPSCs were maintained on feeders in PXGL culture medium: N2B27 (1:1 DMEM/F12:Neurobasal, 0.5X N2 supplement, 0.5X B27 supplement, 1X nonessential amino acids, 2mM L-Glutamine, 1X penicillin-streptomycin, 0.1mM 2-mercaptoethanol, supplemented with 1μM PD0325901 (Stemcell Technologies, #72184), 2μM Gö6983 (Stemcell Technologies, #72462), 2μM XAV939 (Stemcell Technologies, #72674) and 10ng/ml human LIF (Stemcell Technologies, #78055). Medium was refreshed every day for all the cell lines. All the drug treatments were carried out for 48h in S/L media unless mentioned otherwise. PladienolideB (PlaB) treatment: 2.5nM; Curaxin (CBl0137) treatment: 0.5μM; Polyamine inhibition: 0.25mM of DFMO and 15μM of AMXT-1501.

### siRNA transfections

Mouse ESCs were plated in Serum/Lif medium in 6-well plates or in 150mm culture plates at 0.2 x 10^6 cells per well or 4 x 10^6 cells per plate. On the following day, the cells were transfected with a combination of two siRNAs against Zscan4; Ambion: Cat#4390771, ID: s233512 and Horizon discovery: J-162176-02-0050) or ON-TARGETplus Non-targeting Control siRNAs (**Supplementary Table 9**) using Lipofectamine RNAiMAX at 50pM each per well for a 6-well plate and 1nM for 150mm culture plates. The transfection was repeated the following day and cells were harvested 24h post-transfection.

### Fluorescence-activated cell sorting (FACS)

To sort MERVL Tomato-positive and Tomato-negative cells, mouse ESCs were seeded in 150mm culture dishes at 5 million cells per plate. In case of treatments, the compounds were supplemented in the media while seeding. After 48h (or 24h post siRNA transfection), cells were trypsinized and resuspended in PBS. Prior to sorting using The BD FACSAria™ III Cell Sorter, cells were incubated with DRAQ7 (Invitrogen, Cat#D15106) for 10 mins in the dark at room temperature to exclude non-viable cells.

Flow cytometry data was analyzed using FlowJo (v10.10.0).

### Hi-C

Hi-C was performed on FACS-sorted cells for all the experiments. 500,000 cells were fixed with 2% formaldehyde and frozen for at least 12h at -80°C prior to Hi-C experiments. Hi-C was performed using the Arima Hi-C+ kit (Arima Genomics) according to the manufacturer’s instructions. DNA was sheared using Covaris and sequencing libraries were prepared using the Collibri kit (Invitrogen, Cat#A38612024W). For sequencing, PE35 sequencing was carried out using NextSeq 500/550 High Output Kit v2.5 (75 Cycles) (Illumina, Cat# 20024906).

### Hi-C - data analysis

Hi-C data was processed using the nf-core/hic ^76^. Normalization was then carried out using the ICE (iterative correction and eigenvector decomposition) technique from the cooler suite with default parameters ^77^. Hi-C maps were then visualized using HiGlass for further visual analysis ^78^.

### TAD calling

TADs were computed using Chromosight ^46^ detect function (--pattern=borders) on MERVL-negative matrix at 50kb resolution.

### Compartment analysis (A, B and Z)

Compartment calling was done using cooltools eigs-cis function using the GC content as phasing track. A bin was further associated to A compartment if the E1 was positive (and to B if E1 was negative). Log2 ratio matrix was computed with the hicCompareMatrices program from HiCExplorer package. From this matrix we replace NA values by 0 at 200Kbp resolution and we computed the correlation matrix. Then we extracted the PC1 from this correlation matrix. We defined the Z compartment as the top 5% value from the PC1.

### Pile-up and saddle plots

Pile up analysis was done using the Coolpup.py strategy ^79^. All the pile-up analyses are done to study the relationship between contact probability and Zscan4 binding sites. Briefly, an average of intersections between the Hi-C matrix and all the Zscan4 ChIP-seq peaks ^34^, along with flanking regions of 1Mb, was calculated. The probability of contact is depicted by enrichment at the center of the average matrix, often referred to as the pile- up plot. Saddle plots were plotted using Cooltools to visualize genome-wide compartmentalization and contact probability between motif sites (Zscan4, Dux and Z-DNA). For the later, observed/expected HiC contact frequencies were stratified based on the number of motif sites, binned and plotted.

### ChIP-seq data analysis

Published Zscan4 ChIP-seq experiments SRR10487460 and SRR10487465 were reanalyzed ^34^. Briefly, reads were mapped on mm10 using bowtie2. Aligned reads were filtered on MAPQ30 using Samtools. Peaks were called using Macs2 and SRR10487463 as input. Called peaks were then filtered with the IDR procedure at 0.002.

Previous ChIP-seq for histone marks in ESCs and 2CLCs were reanalyzed ^38^. Reads were aligned and filtered the alignments as previously. After that counts matrixes were generated on bins with FeatureCounts from the subread package ^80^. From these counts we used DESeq2 for computing log2 fold change between 2CLC and mESC on bins or to generate normalized counts.

### Immunofluorescence - Z-DNA staining

Mouse ESCs were plated in gelatin-coated µ-Slide 8 Well (IBIDI, #80826) for a day prior to fixation. Fixation was carried out using 4% paraformaldehyde (PFA) for 10 min at room temperature (RT). Cells were then permeabilized using 0.2% Triton X-100 in PBS for 15 min at RT. Protocol adapted from Yin et al was followed for Z-DNA staining ^81^. Briefly, cells were incubated with 1 mg/mL RNaseA (ThermoFisher, #EN0531) and 50 U/mL of RNase III (ThermoFisher, #AM2290) for 1h at 37°C to remove RNA, including Z-RNA. After 3 washes with PBS, blocking was carried out in MAXblock Blocking Medium (Active Motif, #15252) for 1h at 37 °C. Cells were then incubated overnight at 4°C with primary antibodies diluted (1:200) in 3% BSA. After three washes with PBS, cells were incubated with secondary antibodies diluted (1:500) in 1% PBST for 1h at RT. Following two washes with PBS, nuclei were counterstained with DAPI (Invitrogen #D3571) and ProLong™ Glass Antifade Mountant (Invitrogen #36982) was added to each well.

### DNA-FISH probes

The probes were ordered from Arbor Bioscience (MyTags) and were designed to tile the following regions: Probe Pool 1: Chr5: 141600000 - 142200000, labeled with Alexa488 Probe Pool 2: Chr5: 112500000 - 113000000, labeled with Atto647N Probes were prepared according to the manufacturer’s instructions.

### Combined Immunofluorescence and DNA-FISH (Fluorescence in situ hybridization)

Mouse ESCs were plated on gelatin-coated µ-Slide 8 Well (IBIDI, #80826) for a day prior to fixation with 4% paraformaldehyde (PFA) for 10 min at room temperature (RT). Permeabilization was performed on ice for 5 min in 1x PBS containing 0.5% Triton X-100 and 2mM vanadyl-ribonucleoside complex (VRC, NEB #S1402S). Cells were washed with 2X SCC before blocking in 1% BSA prepared in PBS for 30 min at RT. The primary antibody, diluted 1:200 in 3% BSA solution, was added and cells incubated overnight at 4°C. After washes in 2X SCC, the secondary antibody (1:500 dilution in 1% PBST) was added and incubated for 1h at RT. Before proceeding to the DNA-FISH, a second fixation with 2% PFA was performed prior to a second permeabilization with 0.7% Triton X-100 and 2mM VRC. Cells were dehydrated using an ethanol series (85%, 90% and 100% ethanol, 5 min each) before being denatured in preheated 50% formamide in 2xSSC at 80°C for 40 min. After washes with 2X SSC, oligo probes (10pM each probe) diluted in hybridization buffer (10% Dextran sulfate, 2x SSC, 0.5mm EDTA, 0.05% Triton X100, 0.25mg/ml BSA, 0.5mg/ml PVP in water) were incubated with the cells overnight at 45°C. The next day, cells were washed at 45°C 3 times with 50% formamide in 2X SSC and 3 times with 2X SSC. Nuclei were counterstained with DAPI (Invitrogen #D3571) for 2min at RT and ProLong™ Glass Antifade Mountant (Invitrogen #36982) was added to each well.

### Image acquisition

Imaging was performed using Axio Observer.Z1 / 7 (Zeis) inverted with Apotome.2 and alpha Plan-Apochromat objectives (Zeis) x63/1.46 (oil) and x100/1.46 (oil). Image analysis was done on ImageJ 1.54.

### Western Blotting

Cells were harvested and resuspended in 1X Laemmli buffer (Bio-Rad, #161–0747) with 2.5% of β-mercaptoethanol. Proteins were quantified using Pierce 660nm Protein Assay (ThermoFisher Scientific #22662) according to the manufacturer’s instructions. Equal amounts of proteins were loaded, and equilibration of the different samples was examined by Ponceau staining post transfer on nitrocellulose membrane. Antibodies against Zscan4 (Merck Millipore, AB4320), Vinculin (Abcam, ab18058) were used according to standard protocols. The blots were revealed with Odyssey CLX (Licor).

### Bulk RNA-seq

100,000 cells were sorted for both 2CLCs and mouse ESCs for each condition and the RNA was extracted using the TRIzol method. RNA was quantified using Qubit and assessed for quality using BioAnalyzer system (Agilent). The library preparation was done using Library prep stranded total RNA Ribozero Plus Ligation (Illumina) and were sequenced using NextSeq 2000: 10M-50 cyc (SE50) to generate 40 million reads per sample.

### Analysis of bulk RNA-seq data

Reads were aligned on mm10 genome using STAR. Alignments were filtered with Samtools on MAPQ30. Count matrixes for genes or for bins were generated using featuresCount program from the subread suite. For genes we used the vM23 annotation from gencode. Then DESeq2 was used for computing log2 fold change of genes or bins. To identify genomic clusters enriched for up- or down-regulated genes upon treatments in 2CLCs, genome was scanned with a sliding window of 6Mb (step of 2Mb). In each window the number of differentially expressed gene were counted (up- or down- regulated). Within each window the hypergeometric test was used to determine if differentially expressed genes were significantly overrepresented. The pvalues were corrected with the FDR procedure and windows were selected if the adjusted pvalue < 0.05.

### Single-cell RNA-seq (scRNA-seq)

Mouse ESCs and human iPSCs (untreated and treated with curaxin) were subjected to a split-pool combinatorial barcoding method of scRNA-sequencing using single-cell whole-transcriptome kits (Evercode WT Mini v2) from Parse Biosciences according to the manufacturer’s instructions.

### Analysis of scRNA-seq

The ParseBioscience processing pipeline (v1.1.2) was used with default settings to align sequencing reads to the mm10 mouse genome and hg38 human genome. The count matrixes were analyzed with the Seurat R package in order to cluster the data and generate UMAPs.

### DNA Methyl-seq

DNA was extracted and purified from 100,000 FACS-sorted cells for each condition with the Monarch Genomic DNA Purification Kit (NEB Cat# T3010S) following manufacturer’s instructions. DNA was quantified with Qubit. The NEBNext Enzymatic Methyl-seq Kit (New England Biolabs #E7120S) was used to prepare the libraries. 200 ng of DNA was used as input and sheared to 275 bp fragments using Covaris with the following parameters: Duty Factor = 10%, Peak Incident Power = 175 W, Cycles per burst = 200, Duration = 100 s. Fragment size was assessed by a BioAnalyzer system (Agilent Technologies). Manufacturer’s instructions were followed for the subsequent steps. Libraries were quantified using Qubit and size determined by BioAnalyzer. The libraries were sequenced on an Illumina HiSeq4000 sequencer as paired-end 100 base reads following Illumina’s instructions. Image analysis and base calling were performed using RTA 2.7.3 and bcl2fastq 2.17.1.14. The experiment was performed in triplicates.

### Analysis of methyl-seq data

Methyl-seq data were processed with the Bismark ^82^ pipeline using bowtie2 with default parameters. Biological triplicates were merged for downstream analyses. Methylated and unmethylated CG sites were counted within predetermined windows (bin or interval) and the log2 fold change was calculated for 2CLCs vs mouse ESCs.

### Measurement of polyamine concentration

Equal numbers of cells per condition were used to assess concentration of polyamine using the Total Polyamine Assay Kit (Fluorometric) (Sigma MAK349) according to the manufacturer’s instructions.

## Data availability

All raw and processed sequencing data generated in this study have been deposited at the Gene Expression Omnibus (GEO) under accession number GSE291992.

## Acknowledgements

We thank Romain Koszul for assistance with Hi-C experiments and for providing us with the necessary equipment. We thank Sandrine Schmutz, Pierre-Henri Commere and the Flow Cytometry Platform (Institut Pasteur, Paris) for cell sorting, the Biomics Platform (Institut Pasteur, Paris) for Hi-C and scRNA-seq sequencing. Sequencing for Methyl-seq/RNA-seq was performed by the GenomEast Platform, a member of the “France Génomique” consortium (ANR-10-INBS-0009). This work was supported by grants from European Research Council (AdG SUMiDENTITY), Agence Nationale de la Recherche (ANR-19-CE12-0011-01) and the Sjöberg Foundation to A.D.

## Contributions

S.S. and J.-C.C. conceptualized and designed the project, S.S., performed most of the investigations, Y.L.-M., performed most of the bioinformatic analysis, M.S.-L. contributed to DNA-FISH and immunofluorescence experiments, T.T. produced the scRNA-seq data, A.T. provided an optimized Hi-C protocol and assisted S.S. with the initial Hi-C experiments, A.D. contributed to research design, S.S. and J.-C.C. wrote the paper with input from all authors.

