## Supplementary material for "Z-DNA formation induces the totipotent-like state and primes Zscan4-dependent chromatin compartmentalization": Sup Figures

### Extended Data Figure legends

#### Extended data figure 1

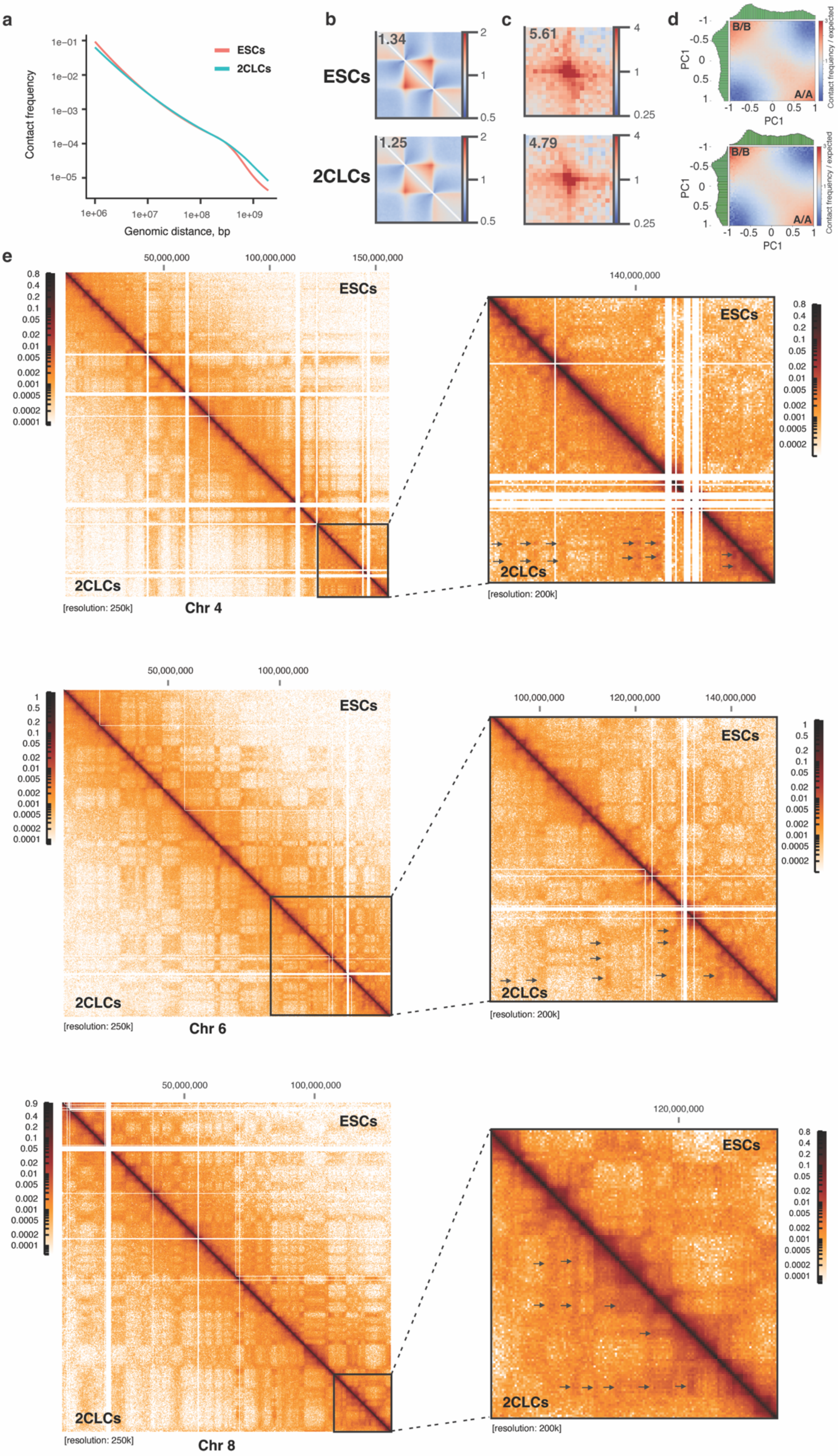

**Extended Data Figure 1. Characterization of the genomic architecture in 2CLCs.**
(**a**) Decay of intra-chromosomal contact probability as a function of genomic distance in ESCs (red) and 2CLCs (cyan). (**b**) Aggregate TAD positions annotated in ESCs and 2CLCS. (**c**) Aggregate over loop positions in ESCs and 2CLCS. (**d**) Saddle plots showing chromatin compartmentalization between 50kb genomic bins ranked according to their eigenvector value (PC1) in ESCs and 2CLCs. (**e**) Example of Hi-C maps for full chromosomes 4, 6, and 8 (left) in ESCs and 2CLCs, along with a zoomed-in view of one end of their subtelomeric regions, opposite the centromere (right).

Extended data figure 2

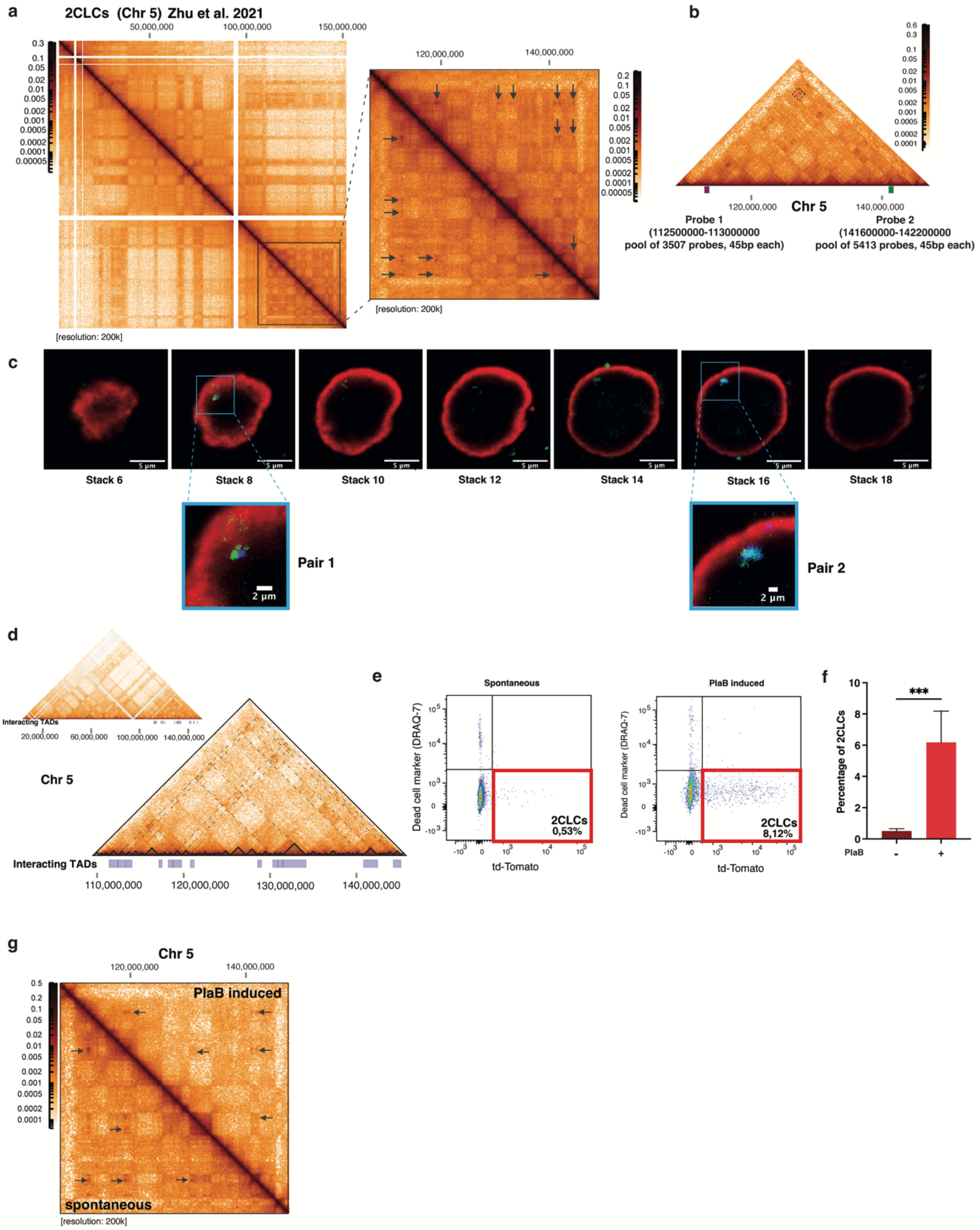


**Extended Data Figure 2. Similar genomic interactions observed in both spontaneous and induced 2CLCs.**

(**a**) Hi-C matrix for chromosome 5 in 2CLCs, published by Zhu et al. (38), with a zoomed-in view highlighting the specific chromatin interactions (arrows) described in Fig. 1a. (**b**) Screenshot showing the positions of DNA-FISH probes 1 (purple) and 2 (green) on chromosome 5, used to visualize the interaction marked by the dotted square on the Hi-C map. (**c**) Representative images obtained as a z-stack (z-resolution = 1 µm) of a Zscan4-

positive 2CLC, showing the overlap of two pairs of DNA-FISH probes. (**d**) Alignment of computed TADs, represented as dark triangles on the Hi-C map of the full chromosome 5 (top), as well as on a zoomed-in region (bottom) in 2CLCs. TADs that show significantly increased interactions in 2CLCs compared to ESCs are represented as a track below the Hi-C map. Dashed lines indicate TAD-TAD interactions. (**e**) Representative dot plots from flow cytometry analysis showing an increase in the MERVL-tdTomato-positive population (2CLCs) upon treatment with Pladienolide B (PlaB) (right) compared to untreated ESCs (left). (**f**) Quantification of the flow cytometry analysis as in “e” (N = 4), *** $p<0.001$, two-tailed t-test. (**g**) Hi-C map illustrating similar 3D genome organization in PlaB-induced 2CLCs (top right) and spontaneous 2CLCs (bottom left) in a defined region of chromosome 5.

Extended data figure 3

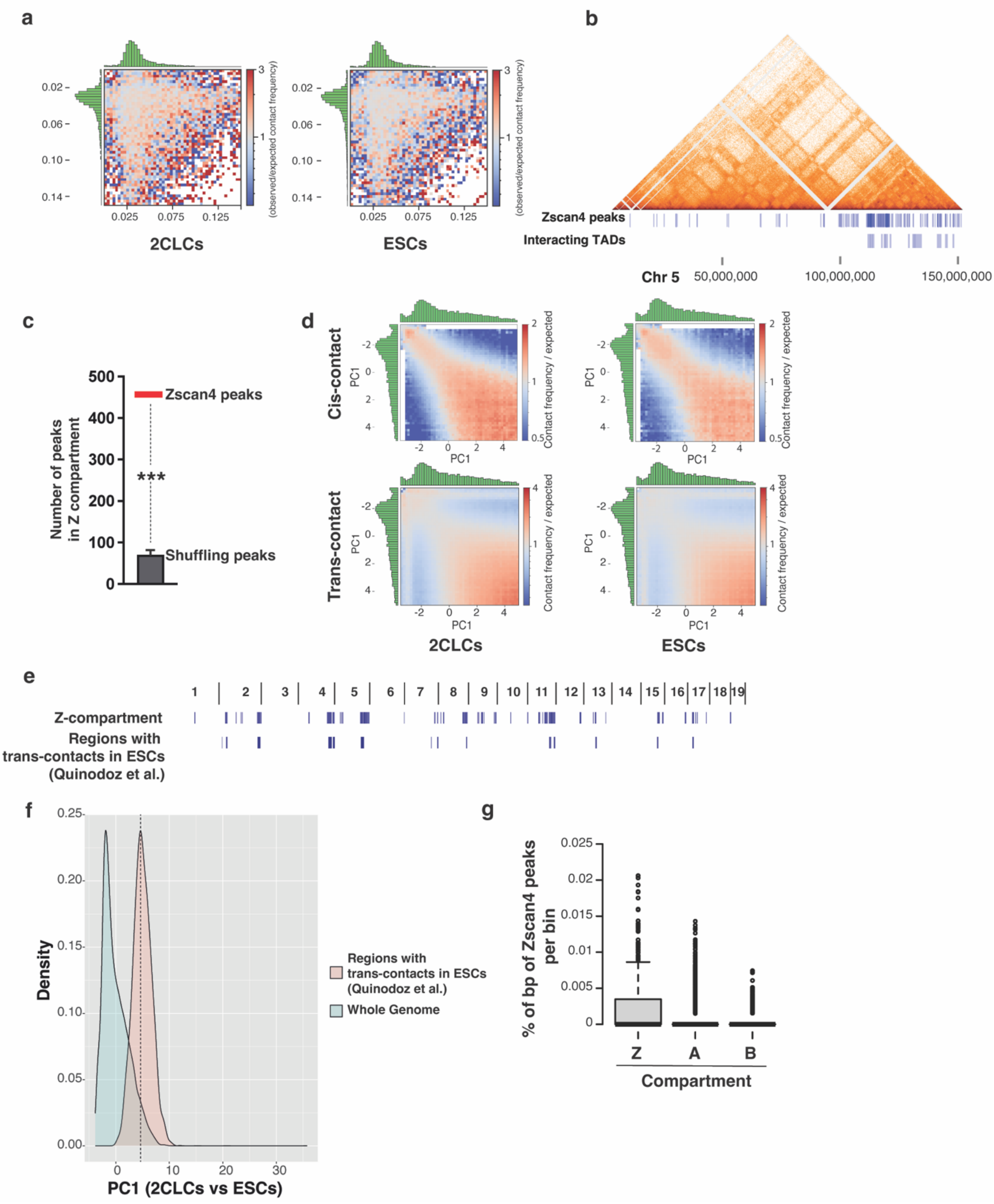


**Extended Data Figure 3. Involvement of Zscan4, but not Dux, in the genomic compartmentalization in 2CLCs.**

(**a**) Saddle plot showing contact probability between 20kb genomic bins ranked according to the number of Dux binding motifs per bin in 2CLCs (left) and ESCs (right). (**b**) Alignment of Zscan4 peaks and 2CLC-specific interacting TADs with the Hi-C map of chromosome 5. (**c**) Number of Zscan4 peaks in Z-compartment (red bar) in comparison to randomly shuffled peaks (1000 sets of random peaks were tested, $p < 1e^{-4}$). (**d**) Saddle plots showing cis- and trans-contact probabilities between 20kb genomic bins, ranked according to their eigenvector value (PC1) from the differential Hi-C analysis between 2CLCs and ESCs. (**e**) Screenshot of the Z-compartment, and the trans-interacting regions identified in ESCs by Quinodoz et al. (55) across the 19 autosomal chromosomes. (**f**) Density distributions of the PC1 value (2CLCs versus ESCs) in regions showing trans-contacts in ESCs from Quinodoz et al. (55) and in whole genome. The dash line represents the PC1 value that has been selected to determine the Z-compartment (Genomic regions within the top 5% of positive PC1 values). (**g**) Percentage of base pairs of Zscan4 peaks per bin of 200kb across the Z, A and B compartment in 2CLCs.

Extended data figure 4

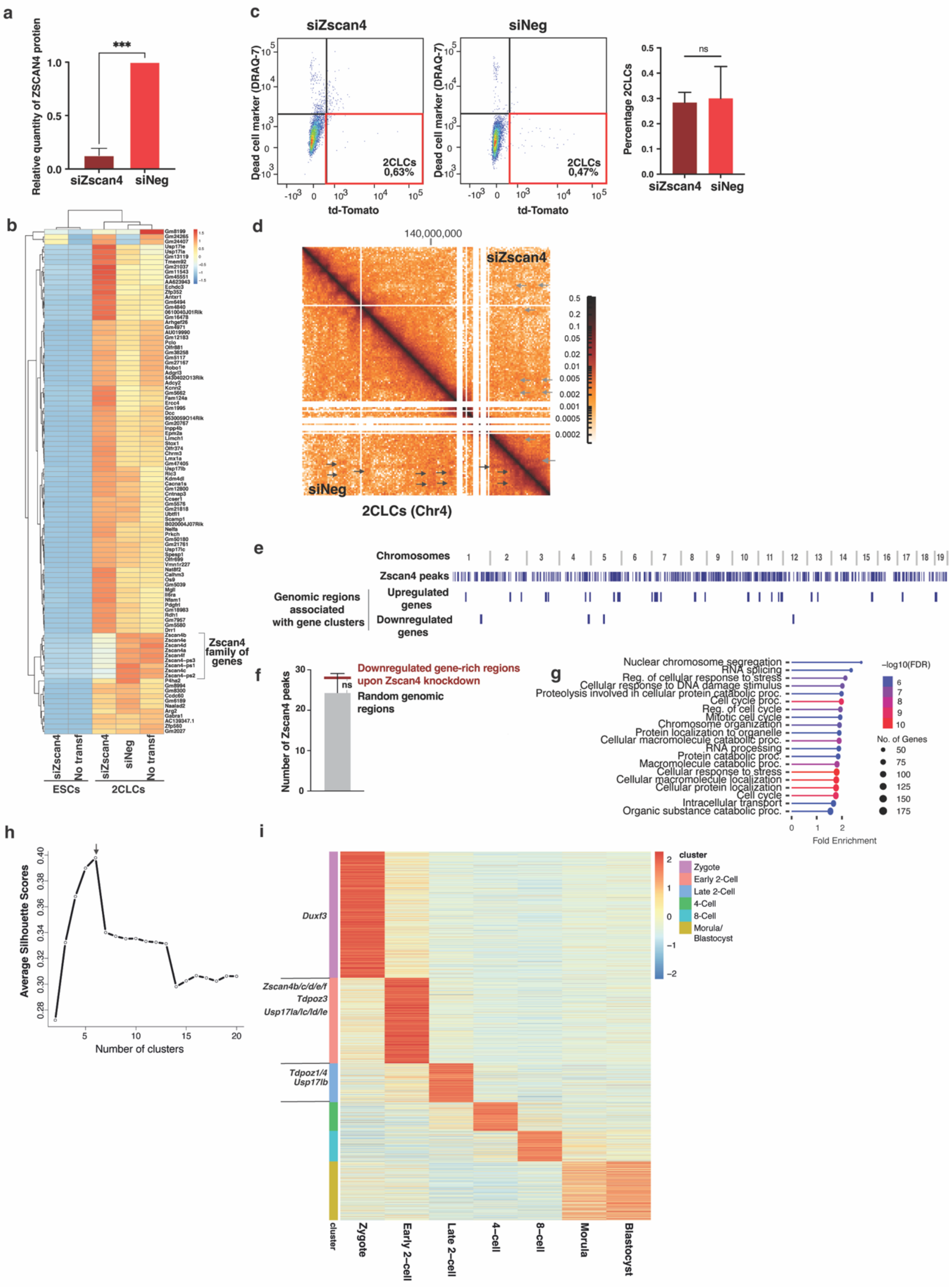


**Extended Data Figure 4. Genomic clustering of upregulated genes upon Zscan4 knockdown in 2CLCs.**

(**a**) Quantification of Zscan4 protein upon knockdown with siRNAs (siZscan4) compared to the non-targeting control (siNeg) in ESCs based on immunoblot analysis, N=4 independent experiments. *** p<0.001, two-tailed t-test. (**b**) Unsupervised hierarchical clustering analysis of the top 100 2CLC markers based on their expression in the indicated samples. The color scale bar represents relative gene expression changes between conditions, normalized by the standard deviation. (**c**) Representative dot plots from flow cytometry analysis after siZscan4 (left) and siNeg (middle) transfection in ESCs. The red square highlights the MERVL-positive 2CLC population. The quantification of N=6 independent experiments is shown on the right. 'ns' indicates non-significant (two-tailed t-test). (**d**) Hi-C map showing 2CLC-specific chromatin interactions on chromosome 4 in control 2CLCs (black arrows) and their reduction in Zscan4-depleted 2CLCs (gray arrows). Resolution: 250kb. (**e**) Localization of autosomal genomic regions enriched for upregulated and downregulated genes in Zscan4-depleted 2CLCs compared to control 2CLCs. The track showing Zscan4 peaks is also indicated. (**f**) Number of Zscan4 peaks in downregulated gene-rich regions upon Zscan4 knockdown in 2CLCs (red bar) in comparison to random regions (1000 sets of random regions were tested), ns for non-significant. (**g**) Ontology analysis of the biological processes associated with downregulated genes upon Zscan4 knockdown in 2CLCs compared to control 2CLCs. (**h**) K-means clustering analysis of genes expressed during early development. The arrow indicates the optimal number of clusters, which is 6. (**i**) Heatmap showing the transcriptional levels of genes in early embryos, consistent with the k-means clustering analysis in panel 'h'.

Extended data figure 5

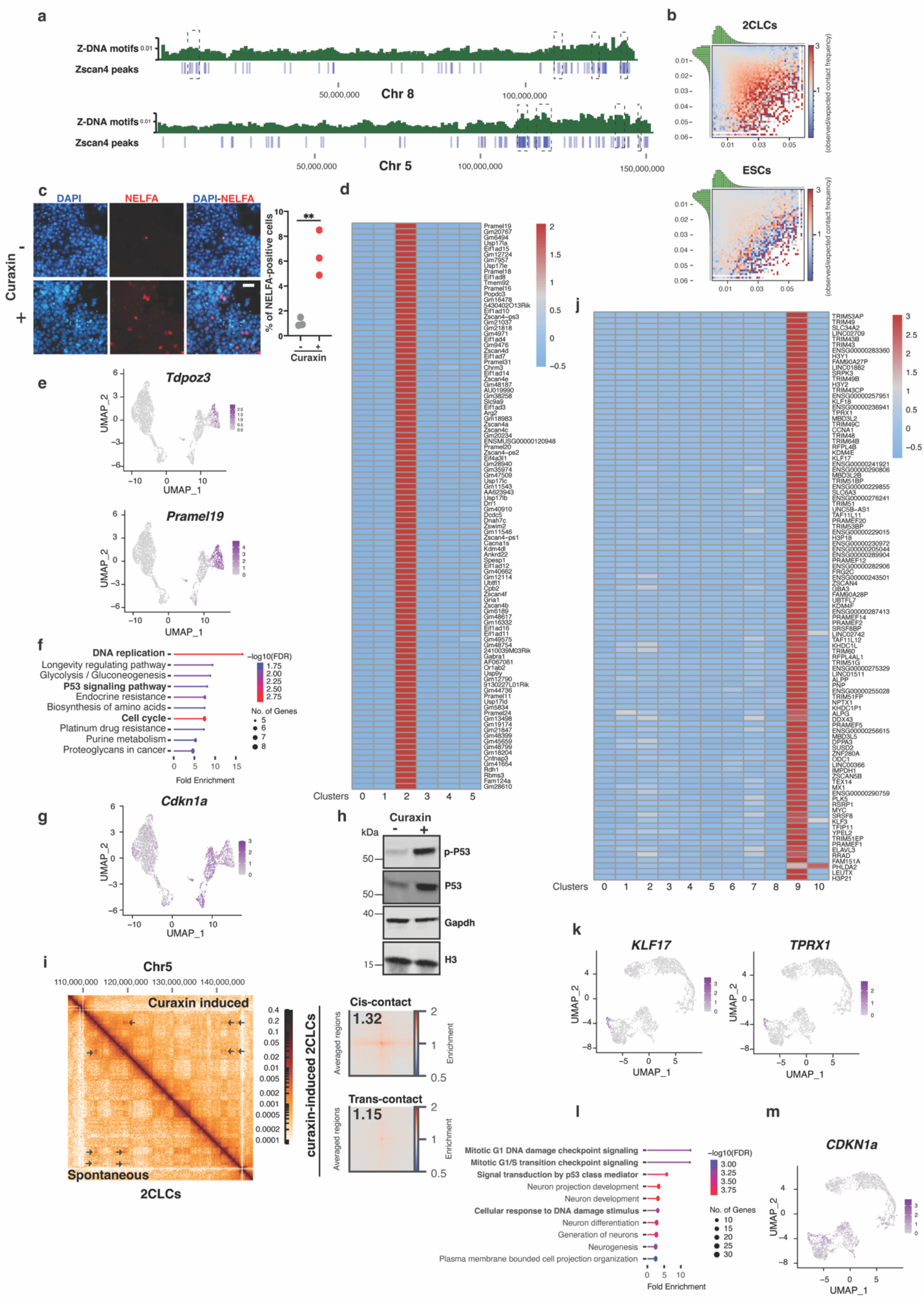


**Extended Data Figure 5. Transcriptomic analysis of the impact of Z-DNA induction on the emergence of murine 2CLCs and human 8CLCs.**

(**a**) Alignment of Z-DNA motifs and Zscan4 peaks along chromosomes 5 and 8. (**b**) Saddle plot showing contact probability between 20kb genomic bins ranked according to the number of Z-DNA motifs per bin in 2CLCs (top) and ESCs (bottom). (**c**) Representative immunofluorescence images showing an increase in the number of 2CLCs stained with an anti-Nelfa antibody (red) in mESCs treated with 0.5 μM curaxin for 48 hours, compared to the control condition, and the associated quantification for N=3 independent experiments, ** $p<0.01$, two-tailed t-test. Scale bar: 50 μm. (**d**) Heatmap showing the relative expression of the top 100 2CLC markers (66) across the six clusters identified from the scRNA-seq analysis of ESCs with or without curaxin treatment (see Fig. 4E). The color scale bar represents relative gene expression changes between conditions, normalized by the standard deviation. (**e**) UMAP plot as in 'Fig. 4e' showing the expression of *Tdpoz3* and *Pramel19*. (**f**) KEGG Pathway enrichment categories for the top 200 gene markers of clusters 1+4 (see Fig. 4E). (**g**) UMAP plot as in 'Fig. 4e' showing the expression of *Cdkn1a*. (**h**) Representative example of immunoblots for P53, phospho-P53 (p-P53) as well as Histone H3 and Gapdh, used as the housekeeping controls in ESCs treated or not with curaxin. (**i**) (Left) contact probability matrix for the indicated portion of the chromosome 5 (Chr 5), comparing the curaxin-induced 2CLCs to spontaneous 2CLCs. (Right) Pile up plots of cis- and trans-contact probability enrichment at Zscan4 binding sites in curaxin-induced 2CLCs. (**j**) Heatmap showing the relative expression of the top 100 human 8CLC markers (66) across the 11 clusters identified from the scRNA-seq analysis of human iPSCs with or without curaxin treatment (see Fig. 4h). The color scale bar represents relative gene expression changes between conditions, normalized by the standard deviation. (**k**) UMAP plot as in 'Fig. 4h' showing the expression of *KLF17* and *TPRX1*. (**l**) KEGG Pathway enrichment categories for the top 200 gene markers of clusters 2+3+4+7+10 (see Fig. 4H). (**m**) UMAP plot as in 'Fig. 4h' showing the expression of *CDKN1a*.

Extended data figure 6

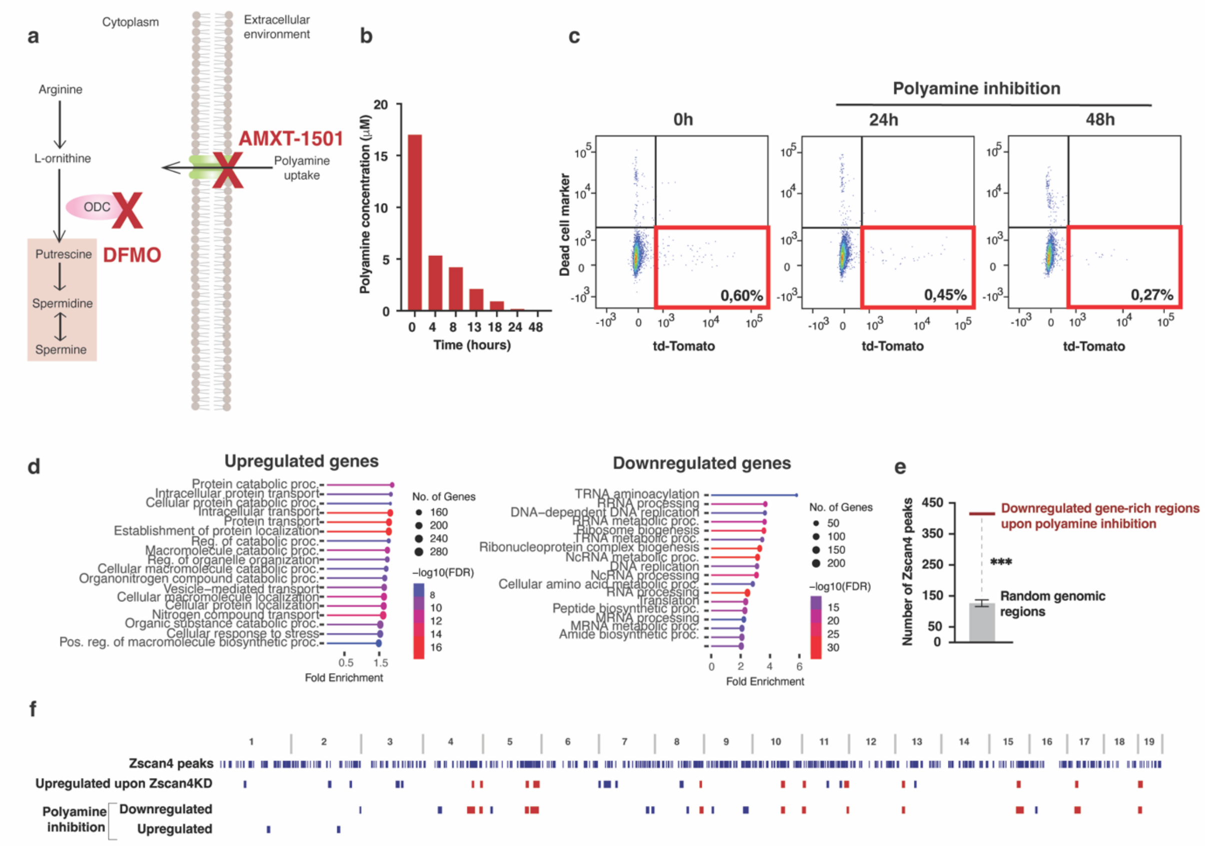


**Extended Data Figure 6. Transcriptomic analysis of 2CLCs upon polyamine inhibition.**

(**a**) A simplified schematic of the polyamine synthesis pathway highlighting the action of DFMO and AMXT-1501 (red crosses) in reducing polyamine concentration in cells. (**b**) Bar graph showing the gradual depletion of polyamine concentration over time in ESCs treated with 0.25 mM DFMO and 15 μM AMXT-1501. N=2. (**c**) Representative dot plots from flow cytometry analysis of control ESCs and ESCs treated for 24 or 48 hours with polyamine pathway inhibitors. The red square highlights the MERVL-positive 2CLC population. (**d**) Ontology analysis of biological processes associated with upregulated genes (left) and downregulated genes (right) in 2CLCs after 48 hours of polyamine inhibition (0.25 mM DFMO and 15 μM AMXT-1501). (**e**) Number of Zscan4 peaks in downregulated gene-rich regions upon polyamine inhibition in 2CLCs (red bar) in comparison to random regions (1000 sets of random regions were tested, $p < 1e^{-4}$). (**f**) Localization of autosomal genomic regions enriched for upregulated and downregulated genes in 2CLCs upon polyamine inhibition compared to control 2CLCs. Tracks displaying Zscan4 peaks and regions enriched for genes upregulated in Zscan4-depleted 2CLCs (as in Extended Data Fig. 4e) are also indicated. Overlapping genomic regions enriched for upregulated genes in Zscan4-depleted 2CLCs and downregulated genes in 2CLCs upon polyamine inhibition are colored in red.

Extended data figure 7

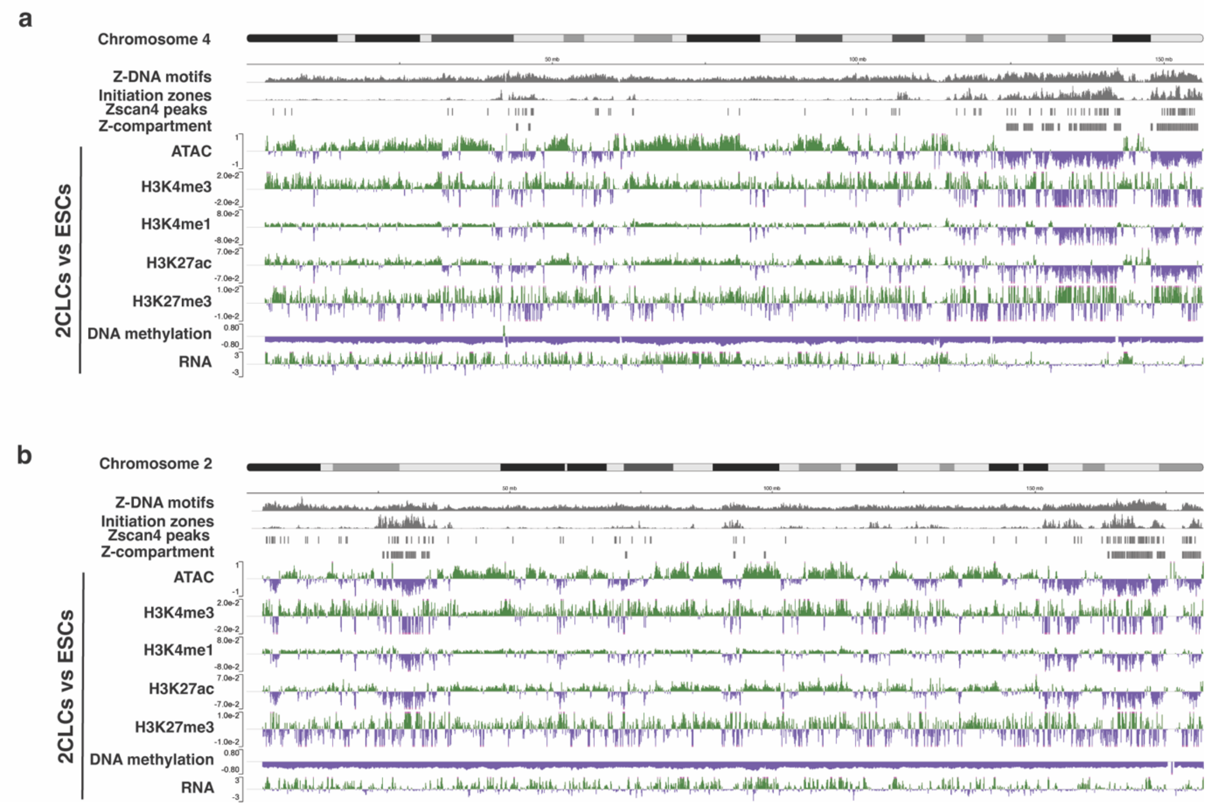


**Extended Data Figure 7. Examples of epigenetic and transcriptomic dynamics during the conversion of ESCs to 2CLCs.**

(**a**) Screenshot of Z-DNA motifs, DNA replication initiation zones, Zscan4 peaks, Z-compartment, and the log2 fold change of histone marks, DNA methylation, and RNA transcription signals between 2CLCs and ESCs across chromosome 4. (**b**) Same as 'a' for chromosome 2.

**Supplementary Table 1**. List of TADs identified in mouse ESCs
**Supplementary Table 2**. List of TAD-TAD interactions identified in 2CLCs
**Supplementary Table 3**. List of Zscan4 peaks in 2CLCs
**Supplementary Table 4**. PC1 values from Hi-C analysis (2CLCs/ESCs) for each bin of 200kb
**Supplementary Table 5**. Log2 Fold changes (2CLCs/ESCs) and means (2CLCs and ESCs) of histone marks, methyl-seq, RNA-seq for each bin of 200kb as well as the coverage of Zscan4 peaks, Z-DNA motifs and DNA initiation zones for each bin of 200kb
**Supplementary Table 6**. Differential analysis of genes and repeats in 2CLCs depleted for Zscan4 compared to control 2CLCs and in DFMO+AMXT-treated 2CLCs compared to control 2CLCs
**Supplementary Table 7**. List of scRNA-seq markers in mouse cells
**Supplementary Table 8**. List of scRNA-seq markers in human cells
**Supplementary Table 9**. Sequence of the siRNAs used in this study
